# Structural insights into CXCR4 modulation and oligomerization

**DOI:** 10.1101/2024.02.09.579708

**Authors:** Kei Saotome, Luke L. McGoldrick, Jo-Hao Ho, Trudy F. Ramlall, Sweta Shah, Michael J. Moore, Jee Hae Kim, Raymond Leidich, William C. Olson, Matthew C. Franklin

**Affiliations:** Regeneron Pharmaceuticals, Inc. Tarrytown, NY 10591

## Abstract

Activation of the chemokine receptor CXCR4 by its chemokine ligand CXCL12 regulates diverse cellular processes. CXCR4 also serves as a key target for diseases such as cancer and HIV. Previously reported crystal structures of CXCR4 bound to antagonists revealed the architecture of an inactive, homodimeric receptor. However, many structural aspects of CXCR4 remain poorly understood, including its activation by CXCL12, as well as its assembly into higher-order oligomers. Here, we use cryoelectron microscopy (cryoEM) to investigate various modes of CXCR4 regulation in the presence and absence of G_i_ protein. CXCL12 activates CXCR4 by inserting its N-terminus deep into the CXCR4 orthosteric pocket. The binding of FDA-approved antagonist AMD3100 is stabilized by electrostatic interactions with acidic residues in the 7 transmembrane helix bundle. A potent antibody blocker, REGN7663, binds across the extracellular face of CXCR4 and inserts its CDR-H3 loop into the orthosteric pocket. Trimeric and tetrameric structures of CXCR4 reveal, to our knowledge, previously undescribed modes of GPCR oligomerization. Remarkably, CXCR4 adopts distinct subunit conformations in trimeric and tetrameric assemblies, highlighting how oligomerization could allosterically regulate chemokine receptor function.

## Main

Chemokine receptors are a family of Class A G-protein coupled receptors (GPCRs) that mediate cell migration in response to binding of chemokine ligands^1^. CXCR4 is a well-studied chemokine receptor that is activated by the chemokine ligand CXCL12 (also called stromal cell-derived factor 1, or SDF-1) and signals primarily through coupling with G_i_ protein^2^, regulating cell migration in hematopoeisis, neovascularization, angiogenesis and various other physiological processes^3^. CXCR4 is involved in numerous diseases, including roles as a cancer marker implicated in tumor proliferation^4^ and as a coreceptor for X4-tropic HIV strains^5^. Mutations in CXCR4 that result in enhanced and prolonged signaling result in a rare immune disorder called WHIM (warts, hypogammaglobulinemia, infections, and myelokathexis) syndrome^6^. The significant roles of CXCR4 in health and disease have made the receptor an intensely investigated drug target^7^. The small molecule CXCR4 antagonist AMD3100 (plerixafor), initially developed as an HIV entry inhibitor^8^, was FDA-approved as a hematopoietic stem cell mobilizer for autologous transplantation in patients with Non-Hodgkin’s lymphoma or multiple myeloma^9,10^. Numerous additional CXCR4-targeting therapeutics have been developed^7^, notably including monoclonal antibodies with improved pharmacokinetic properties and thus potentially greater efficacy compared to small molecules and peptides^11–13^.

Structural studies of Class A GPCRs have focused on isolated monomeric forms of the receptors bound to various ligands, pharmacological modulators, and transducer proteins^14^. However, increasing evidence suggests GPCRs can form dimers and higher order oligomers in the plasma membrane, with implications for signaling and therapeutic action^15^. Chemokine receptors are no exception; a multitude of studies have indicated the existence of homo- and hetero-oligomers^16–18^, including crystal structures of antagonist-bound CXCR4 consistently revealing homodimeric forms^19,20^. Interestingly, CXCR4 has also shown a propensity to form higher order oligomers using a mechanism that can be separated from dimerization^21^.

Despite its critical roles in health and disease, many mechanistic aspects of CXCR4 remain poorly understood, owing to a lack of structural information. These include its activation by CXCL12, binding mode of AMD3100, coupling to G_i_ protein, inhibitory action of antibodies, and mechanisms of higher order oligomerization. Here, we address these open questions by reporting a series of cryoelectron microscopy (cryoEM) structures of CXCR4 complexes.

## Results

### Structural basis of CXCL12 and AMD3100 action on CXCR4

To stabilize active state signaling complexes and improve protein yields we made the following modifications to wild type CXCR4: we replaced the N-terminal methionine with an HA (hemagglutinin) signal peptide^22^, included a previously characterized constitutively active mutation (N119S)^23^, and fused monomeric eGFP^24^ and FLAG tag to the receptor C-terminus (Extended Data Fig. 1a). We refer to this construct as CXCR4_EM_. We also employed a Gα_i_ construct harboring dominant negative mutations^25^ to facilitate isolation of receptor/G_i_ complexes in the absence of stabilizing antibody fragments^26^. Fluorescence detection size exclusion chromatography (FSEC)^27^ experiments indeed indicated complex formation between CXCR4_EM_ and G_i_ in the absence of agonist (Extended Data Fig. 1b). We prepared detergent-solubilized CXCR4_EM_/G_i_ complexes and first determined cryoEM structures in apo, CXCL12-bound, and AMD3100-bound states at overall resolutions of 2.7, 3.3, and 3.2 Å, respectively (Fig. 1, Extended Data Figs. 2, 3). Each of the structures shows a prototypical arrangement of an active receptor coupled to heterotrimeric G protein, including a hallmark kink of TM6 relative to previously reported crystal structures of antagonist-bound CXCR4^19,20^ (Extended Data Fig. 4a). We therefore refer to the CXCR4 conformation in these structures as active.

**Figure 1.**
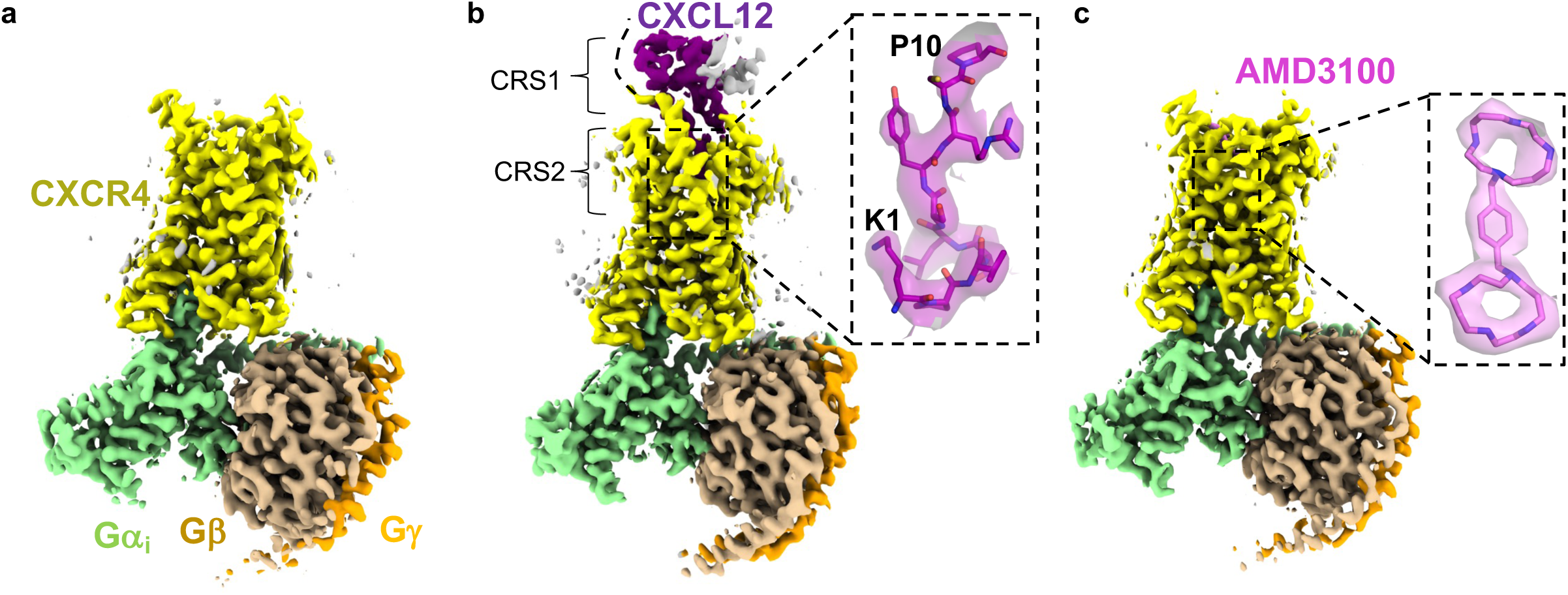
CryoEM reconstructions of CXCR4/G_i_ complexes. **a**, apo CXCR4/G_i_ complex. **b,** CXCR4/G_i_/CXCL12 complex. Inset shows fit of CXCL12 N-terminal tail (res. 1-10) in cryoEM map, shown as semitransparent surface. Locations of chemokine recognition sites 1 and 2 are labeled. Curved dotted line represents missing density for distal N-terminus of CXCR4, which has been reported to interact with CXCL12. **c**, CXCR4/G_i_/AMD3100 complex. Inset shows fit of AMD3100 compound in cryoEM map.

Our cryoEM reconstruction of CXCR4_EM_/G_i_/CXCL12 revealed clear signal for the chemokine bound at the extracellular side of the receptor (Fig. 1a). Density for the chemokine N-terminus (res. 1-12) was sufficiently resolved to build side chains (Fig. 1b), whereas the remainder of the ligand was less resolved due to flexibility and only permitted main chain tracing (Extended Data Fig. 3i). Consequently, interactions between the chemokine N-terminal region and receptor orthosteric pocket (chemokine recognition site 2) were readily discernible, while interactions between the globular portion of the ligand and the N-terminus of CXCR4^28^ (chemokine recognition site 1) were unclear. CXCL12 is known to exist in monomeric and dimeric forms that have been shown to yield distinct signaling outcomes upon CXCR4 binding^29,30^. Weak signal corresponding to a second protomer of the CXCL12 dimer could be observed in our cryoEM reconstruction, consistent with the notion that dimeric forms of CXC ligands act on single receptor subunits^20,31^ (Extended Data Fig 3i).

The binding mode of CXCL12 onto CXCR4 is overall similar to those found in published structures of CC and CXC chemokine/chemokine receptor complexes^31–36^ (Fig. 2, Extended Data Fig. 4b). However, the CXCL12 binding pose observed in our structure notably differs from that of CXCL12 bound to atypical chemokine receptor 3 (ACKR3, formerly known as CXCR7)^37^, a promiscuous receptor that has been suggested to function as a chemokine “scavenger” and has approximately 10-fold higher affinity for CXCL12 than CXCR4^38,39^. The CXCL12 C-terminal α helix is rotated ∼70° when bound to ACKR3 relative to CXCR4 (Extended Data Fig. 4c). Correspondingly, the 40s loop of CXCL12 is situated proximal to the N-terminal region in CXCR4, while it is nearby ECL3 in ACKR3. In addition to the distinct overall chemokine/receptor docking orientations, the binding geometries of the CXCL12 N-terminus within the orthosteric pockets of each receptor are also unique (Extended Data Fig. 4d).

**Figure 2.**
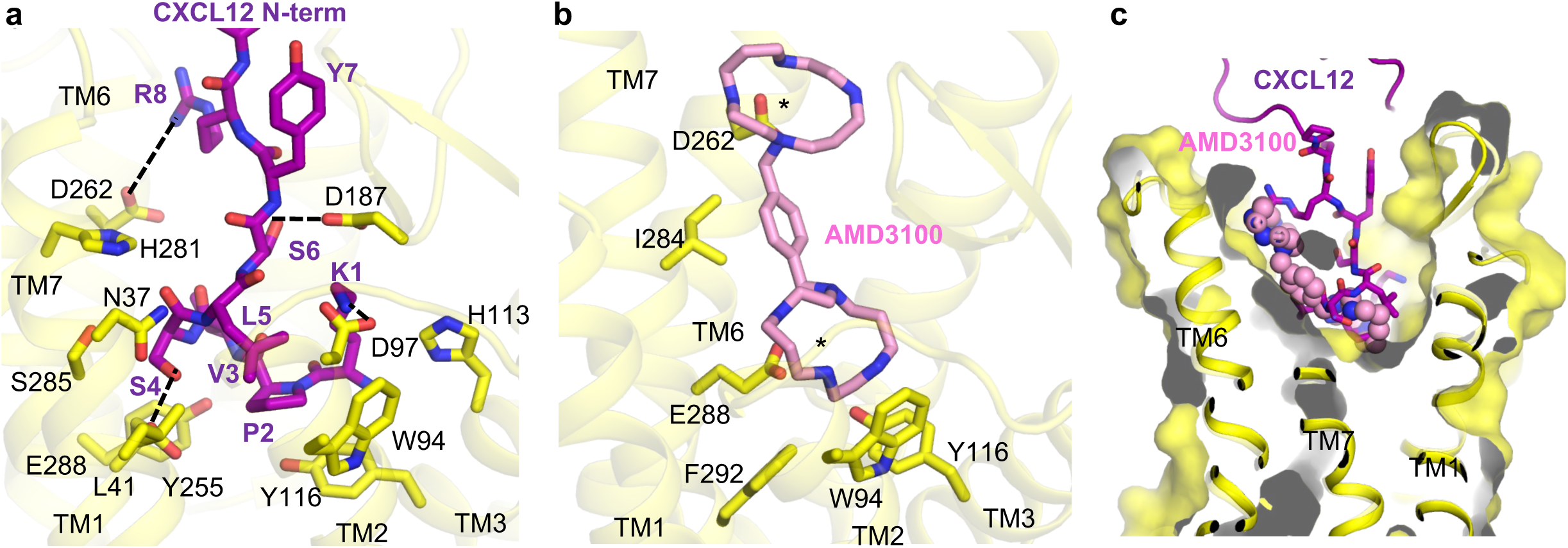
Interactions between CXCR4 and ligands. **a**, expanded view of interaction between CXCL12 N-terminal tail and CXCR4 orthosteric pocket. Hydrogen-bonding and electrostatic interactions are depicted as dashed lines. **b,** expanded view of AMD3100 binding at CXCR4 orthosteric pocket. Asterisks indicate positions of the two lactam rings, each of which interact with acidic residues. **c,** cutaway surface view of CXCR4 orthosteric pocket. CXCL12 N-term is shown as sticks and AMD3100 is shown as spheres to illustrate their relative binding positions in the orthosteric pocket.

Mutations at the distal N-terminus of CXCL12 can convert the chemokine into an antagonist^40^, highlighting its importance for receptor activation. Our structure shows how the CXCL12 N-terminus protrudes into the orthosteric pocket of CXCR4 and makes extensive contacts with the TM core (Fig. 2a). The distal CXCL12 N-terminus is positioned overall deeper into the pocket than that of the antagonistic viral chemokine vMIP-II^20^ (Extended Data Fig. 4e), consistent with their respective ligand functions. P2_CXCL12_ penetrates deepest into the orthosteric pocket, contacting the side chain of Y116^3.32^ (Ballesteros-Weinstein numbering^41^ in superscript). The side chain of K1_CXCL12_ projects upward from the TM core to the extracellular side of the receptor and is positioned to interact electrostatically with D97^2.63^ and possibly D187^ECL^^2^. S4_CXCL12_ makes an apparent hydrogen bond interaction with E288^7.39^. L5_CXCL12_ packs onto a mainly hydrophobic surface composed of L41^1.35^, Y45^1.39^, W94^2.60^, A98^2.64^. R8_CXCL12_ appears poised to make a charge-charge interaction with D262^6.58^, as predicted previously based on charge-swap experiments^28^. Several of the CXCR4 residues mentioned above (W94^2.60^, D97^2.63^, Y116^3.32^, D187^ECL2^, E288^7.39^) have been shown to be important for CXCL12/CXCR4 signaling^28,42,43^, underscoring the functional relevance of the interactions observed in our cryoEM structure. We expand on the structural basis of CXCL12 activation of CXCR4 in a following section.

We observed unambiguous density for the bilobed AMD3100 molecule in our cryoEM reconstruction of CXCR4_EM_/G_i_/AMD3100 (Fig. 1c). Although it has primarily been described as an antagonist^44^, our observation that AMD3100 binds to the active CXCR4_EM_/G_i_ complex without disrupting G protein coupling is consistent with the compound acting as a weak partial agonist on constitutively active mutants of CXCR4. AMD3100 binds the orthosteric pocket using a diagonal orientation and directly blocks CXCL12 docking, although its overall binding mode is shifted toward TM5/6 relative to the CXCL12 N-terminus (Fig. 2b,c). Each of the two positively charged cyclam rings of AMD3100^45^ is stabilized electrostatically by an acidic side chain pointed toward the center of the ring; the cyclam moiety closer to the extracellular side interacts with D262^6.58^ while the cyclam proximal to the transmembrane core interacts with E288^7.39^. The closely matched spacings between the side chains of D262^6.58^ and E288^7.39^ residues and the cyclam rings therefore appears be the main binding determinant of AMD3100 and other bicyclam analogues. Consistent with our structure, a previous study showed that that D262N and E288A mutants each reduced the affinity of AMD3100 to CXCR4 by more than 50-fold^45^. The central aromatic ring of the phenylenebis(methylene) linker connecting the two cyclam moieties makes hydrophobic contacts with I284^7.35^, which is positioned directly in between D262^6.58^ and E288^7.39^ in the orthosteric pocket. This interaction may contribute to the increased potency of bicyclams with an aromatic linker rather relative to those with an aliphatic linker^46^. The rigidity imposed by the aromatic linker on the relative positions of the two cyclam moieties may also play a role in stabilizing the binding pose of AMD3100.

### Antagonism of CXCR4 by REGN7663 mAb

Antibody-based therapeutics against CXCR4 and other GPCRs are a promising alternative to small molecules due to their high specificity to the target, opportunity for Fc-effector functions, and favorable pharmacokinetic properties^11,13,47,48^. REGN7663 is a fully human anti-CXCR4 monoclonal antibody (mAb) generated using VelocImmune mice^49,50^. We showed using a CRE-Luciferase reporter assay that REGN7663 is a potent blocker (IC50=2.72 nM) of CXCL12-induced signaling in HEK293 cells engineered to overexpress CXCR4 (Fig. 3a). Further, in the absence of CXCL12, REGN7663 decreased the apparent basal activity (EC50=1.71 nM), indicating inverse agonism in the setting of CXCR4 overexpression (Fig. 3b). To understand how REGN7663 binds and inhibits CXCR4, we determined a 3.4 Å resolution cryoEM structure of REGN7663 Fab in complex with CXCR4_EM_/G_i_ (Fig. 3c, Extended Data Fig. 5a-d). The structure revealed that REGN7663 binds directly onto the extracellular face of CXCR4, antagonizing the receptor by steric blockade of CXCL12 binding. Most of the REGN7663 epitope resides at the extracellular N-terminal region and ECL2 (Extended Data Fig. 5e,f). The REGN7663 heavy chain dominates the binding interactions, burying significantly more surface area (∼1100 Å^2^) than the light chain (∼300 Å^2^). Although the overall architecture of the complex is similar to the apo, CXCL12-bound, and AMD3100-bound CXCR4_EM_/G_i_ structures, REGN7663 binding induces distinct conformations of the N-terminus and ECL2, suggesting their flexibility is important for specific mAb binding (Extended Data Fig. 5g). Heavy chain complementarity-determining regions (CDRs) 1 and 2 of REGN7663 are oriented toward the extracellular ends of TM4 and TM5, while light chain CDRs are oriented extracellular to TM1 and TM2 (Fig. 3d). Remarkably, the CDR-H3 loop of REGN7663 wedges between the CXCR4 N-terminus and ECL2, exhibiting a partial insertion into the CXCR4 orthosteric pocket. The side chain of REGN7663 residue R105 protrudes deepest into the orthosteric pocket, making an apparent charge-charge interaction with E288^7.39^(Fig. 3e). The insertion of CDR-H3 loop, though not activating in the case of REGN7663, is reminiscent of how the CDR3 loop of the single domain antagonist antibody JN241 occupies the orthosteric pocket of apelin receptor^51^. Taken together with the finding that JN241 was converted into a full agonist through subtle engineering of CDR3^51^, our structure of REGN7663/CXCR4 complex illustrates the potential for full antibodies (containing light and heavy chains) functionally modulating GPCRs by inserting CDR loop(s) into the orthosteric pocket.

**Figure 3.**
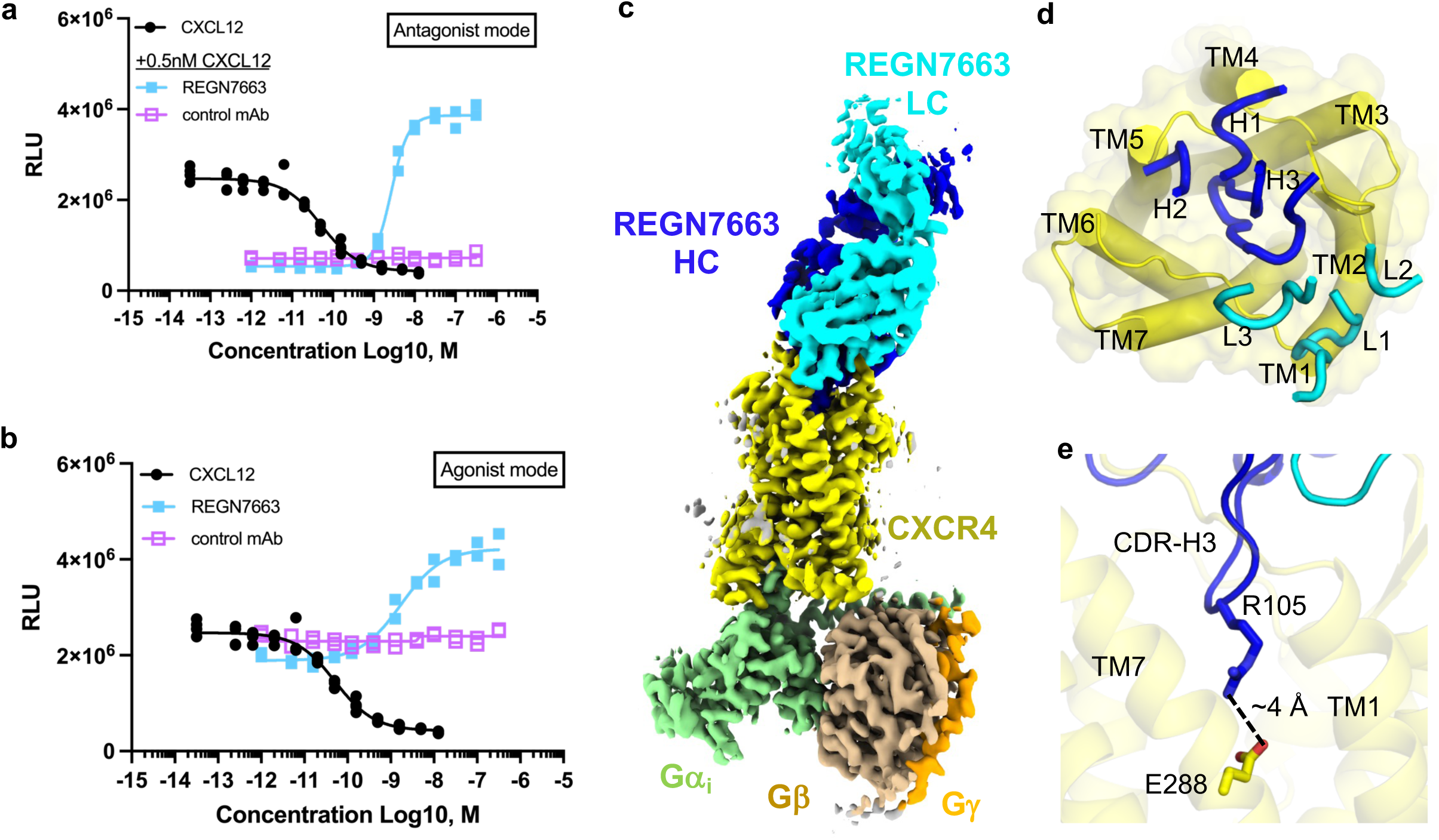
CXCR4 antagonism by REGN7663 mAb. **a**, CRE-Luciferase reporter assay showing CXCL12-dependent decrease in signal and block of CXCL12 activity (at 0.5 nM CXCL12) by REGN7663. **b,** REGN7663 shows concentration-dependent increase in signal relative to baseline in the absence of CXCL12, demonstrating inverse agonism. In (a) and (b), data from two replicate experiments are shown for the REGN7663 and control mAb, and four replicates are shown for CXCL12 (the same data for CXCL12 are shown in (a) and (b) to allow comparison with mAb data). **c,** cryoEM reconstruction of CXCR4_EM_/G_i_/REGN7663 Fab complex, with each polypeptide chain colored differently. **d,** top-down view of CXCR4 (yellow) with CDR loops of bound REGN7663 shown (blue = heavy chain, cyan = light chain). **e,** electrostatic interaction between CDR-H3 of REGN7663 and CXCR4 orthosteric pocket-facing residue E288.

### Conformational changes associated with CXCR4 activation and Gα_I_ protein docking

We next sought to assess the conformational changes associated with CXCR4 activation. Available crystal structures of inactive, antagonist-bound CXCR4 contain construct modifications, namely T4 lysozyme (T4L) inserted at ICL3 and a thermostabilizing mutation in TM3, that could confound comparison with our current structures. We therefore determined a 3.1 Å resolution cryoEM structure CXCR4_EM_ in the absence of G_i_ protein, utilizing REGN7663 Fab as a fiducial mark (Fig. 4a, Extended Data Fig. 5h-k). Structural alignment of the REGN7663 Fab/CXCR4_EM_/G_i_ structure with the G_i_-free REGN7663 Fab/CXCR4_EM_ structure showed nearly identical conformations at the REGN7663 epitope/paratope regions but distinct conformations at the intracellular half of the receptor, including the characteristic movement of TM6 underlying receptor activation (Fig. 4b, Extended Data Fig. 5l). Additional conformational changes upon activation/G_i_-binding include movement of TM5 toward TM6, subtle displacement of TM2 outward, an inward kink of TM7, and loss of ordered structure in H8. We note that H8 was also unresolved in previously determined antagonist-bound CXCR4 crystal structures^19,20^, suggesting this is a consistent feature of the inactive receptor.

**Figure 4.**
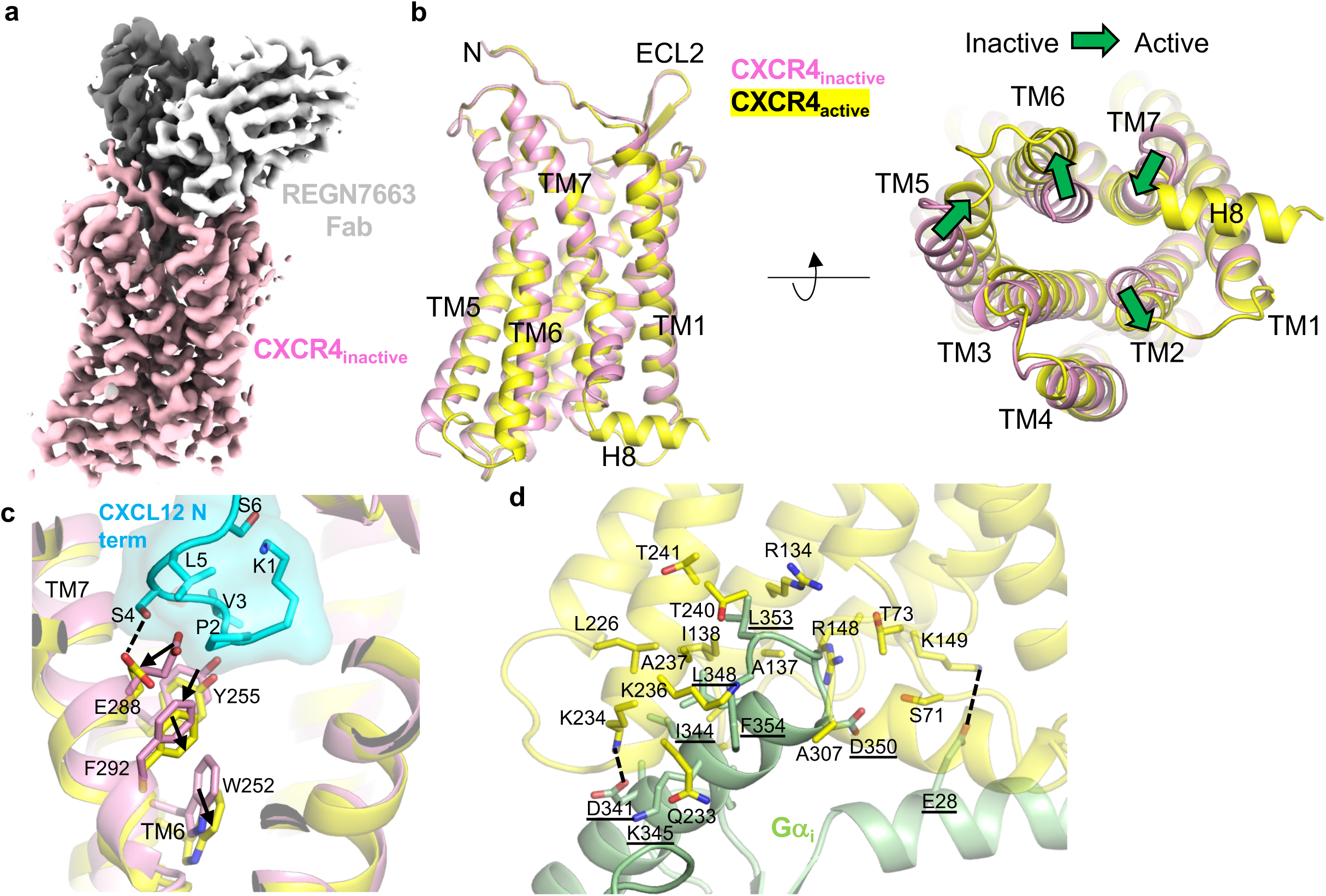
Inactive CXCR4 structure and structural bases of activation. **a**, cryoEM reconstruction of inactive CXCR4_EM_/REGN7663 Fab complex (CXCR4=pink, REGN7663 heavy chain=gray, REGN7663 light chain=white). **b,** structural alignment of inactive CXCR4 (pink) and active CXCR4 (yellow, CXCR4_EM_/G_i_/REGN7663 Fab complex was used for alignment). Side view is shown on left and bottom-up view is shown on right. Green block arrows depict conformational transitions from inactive to active CXCR4. **c,** expanded view showing CXCL12 N-term (cyan) binding to active CXCR4 (yellow). Inactive CXCR4 (pink) is shown for comparison and residues important for transmitting chemokine binding into activation are shown as sticks. **d,** expanded view of Gα_i_ (light green) binding to active CXCR4 (yellow). Residues participating in interaction are shown as sticks and labeled (Gα_i_ residue labels underlined). Electrostatic interactions are highlighted with dashed lines.

We further compared the conformations of the inactive and CXCL12-bound structures to analyze how CXCL12 binding results in activation (Fig. 4c). Binding of CXCL12 N-terminal coil to the orthosteric pocket requires structural changes to the inactive state pocket. Residues P2 and S4 at the CXCL12 N-terminus push E288^7.39^ outward and toward the cytoplasmic side, while V3_CXCL12_ forces a downward displacement of Y255^6.51^. The movements of E288^7.39^ and Y255^6.51^ are in turn transmitted to F292^7.43^, which has been previously implicated in CXCR4 signal transmission^43^, and conserved toggle switch residue^52^ W252^6.48^, respectively. Together, these conformational changes trigger further structural rearrangements that ultimately stabilize the active, G_i_-bound conformation of CXCR4. Furthermore, a close comparison revealed that due to binding of the CXCL12 N-terminus in the orthosteric pocket, E288^7.39^ side chain reorients, along with a small, ∼0.7-1 Å outward movement of the extracellular half of TM7 helix relative to our AMD3100/CXCR4_EM_/G_i_, REGN7663 Fab/CXCR4_EM_/G_i_, and apo CXCR4_EM_/G_i_ structures (Extended Data Fig 6a). This slight conformational difference at TM7 induced by CXCL12 may explain why it is full agonist, while the other ligands are not. Similar structural mechanisms of chemokine activation to that described above for CXCL12 have been observed for the CCR2/CCL2 complex^32^ and CCR5/MIP-1α complex^34^.

Like other class A GPCRs, coupling of Gα_i_ to CXCR4 is mediated by insertion of the Gα_i_ α5 helix and C-terminal “wavy hook” into the cytoplasmic-facing core of the receptor TM domain (Fig. 4d). “Wavy hook” residues L353 and F354 bury deepest into CXCR4 and contact R134^3.50^, Q233^ICL3^, K236^6.32^, A237^6.33^, T240^6.36^, and A307 mainly via van der Waals and hydrophobic interactions. Gα_i_ α5 helix makes numerous additional contacts with TM2, TM3, ICL2, TM5, ICL3, and TM6. Salt bridge interactions between D341(Gα_i_)/K234^6.30^ and E28(Gα_i_)/K149 probably play an important role in stabilizing the docking of G_i_ protein onto CXCR4. Although the overall G_i_ binding mode of CXCR4 and other chemokine receptors is shared, the angle at which the Gα_i_ α5 helix docks into the TM bundle differs slightly (Extended Data Fig. 6b). While CXCR4, CXCR1^36^, and CXCR2^31^ show highly similar α5 docking angles, the docking angles in CCR1^33^, CCR2^32^, and CCR5^34^ are similar to each other and shifted relative to CXCR4, owing to distinct intracellular loop conformations and receptor interactions with Gα_i_ (Extended Data Fig. 6c). More specifically, in the CC chemokine receptors, Gα_i_ α5 helix is shifted toward ICL2 and further from ICL3. Available data therefore indicate that CXC and CC chemokine receptors have slightly different G_i_ docking geometries.

### Oligomeric structures of CXCR4

Although GPCRs are generally understood to function as monomeric units, numerous studies have reported that chemokine receptors form dimers and higher order oligomers at the cell surface as expression levels increase^53–56^. Homo- and hetero-oligomerization have been proposed to add complexity to chemokine receptor function, perhaps through allosteric communication between interacting subunits^57,58^. Multiple structures of CXCR4 from different crystal forms showed the same homodimeric architecture^19,20^, demonstrating that the detergent-solubilized receptor has the propensity to dimerize using specific intersubunit interactions mainly involving TM5 and TM6. Our size exclusion chromatography (SEC) data of CXCR4_EM_ consistently showed multiple peaks with different elution volumes, including peaks corresponding to oligomeric species larger than monomeric CXCR4_EM_ or CXCR4_EM_/G_i_ (Extended Data Fig. 1b, 2a). Wild type CXCR4 fused to GFP showed a similar FSEC profile to CXCR4_EM_, indicating that the apparent oligomerization was not specific to the constitutively active N119S mutation present in CXCR4_EM_. We isolated and characterized a presumed oligomeric SEC peak (Extended Data Fig. 2a) of CXCR4_EM_ using cryoEM. Initial cryoEM data yielded clear top/bottom views of trimeric and tetrameric species, but preferred orientation precluded structure determination. After screening various sample preparation conditions, we ultimately employed stage-tilted data collection^59^ to obtain 3.4 Å resolution reconstructions of CXCR4_EM_ homotrimers and homotetramers in complex with REGN7663 Fab (Fig. 5, Extended Data Fig. 7a-j). According to 3D classification, our data contained a roughly 1 to 3 ratio of trimers to tetramer particles (Extended Data Fig. 7k). We did not observe 2D or 3D class averages consistent with dimeric CXCR4, excepting non-physiological antiparallel dimers in our samples prepared in the presence of G_i_ (Extended Data Fig. 7i). The trimer and tetramer both show CXCR4 subunits arranged symmetrically around a cavity at the central axis, at first glance evoking structural similarity to homomeric ion channels, though CXCR4 has no known channel function. In the case of the CXCR4 oligomers, we found evidence for numerous bound lipids at the central axis in the cryoEM maps (Fig. 5 c,f, Extended Data Fig. 8). Due to matching shape features, we tentatively built three phosphatidic acids and three cholesterol molecules in the trimeric map central cavity and four phosphatidic acids and eight cholesterols in the tetrameric cavity (Extended Data Fig. 8d,h). Although the presumed cholesterol molecules could in principle correspond to exogenously added cholesteryl hemisuccinate present in the purification buffers, the EM density we have modeled as phosphatidic acid strongly resembles a phospholipid, and not the LMNG detergent used for purification. This implies that the central cavity lipids were carried over from the cell membrane and remained stably bound through purification, indicating that the oligomeric structures reported here are representative of species present in the CXCR4-expressing cells used in this study and not an artifact of the purification process. The presence of ordered lipids plugging the central axis of CXCR4 oligomers is reminiscent of microbial channelrhodopsin trimers, though the quarternary arrangement of the 7-TM protomers differs^60,61^.

**Figure 5.**
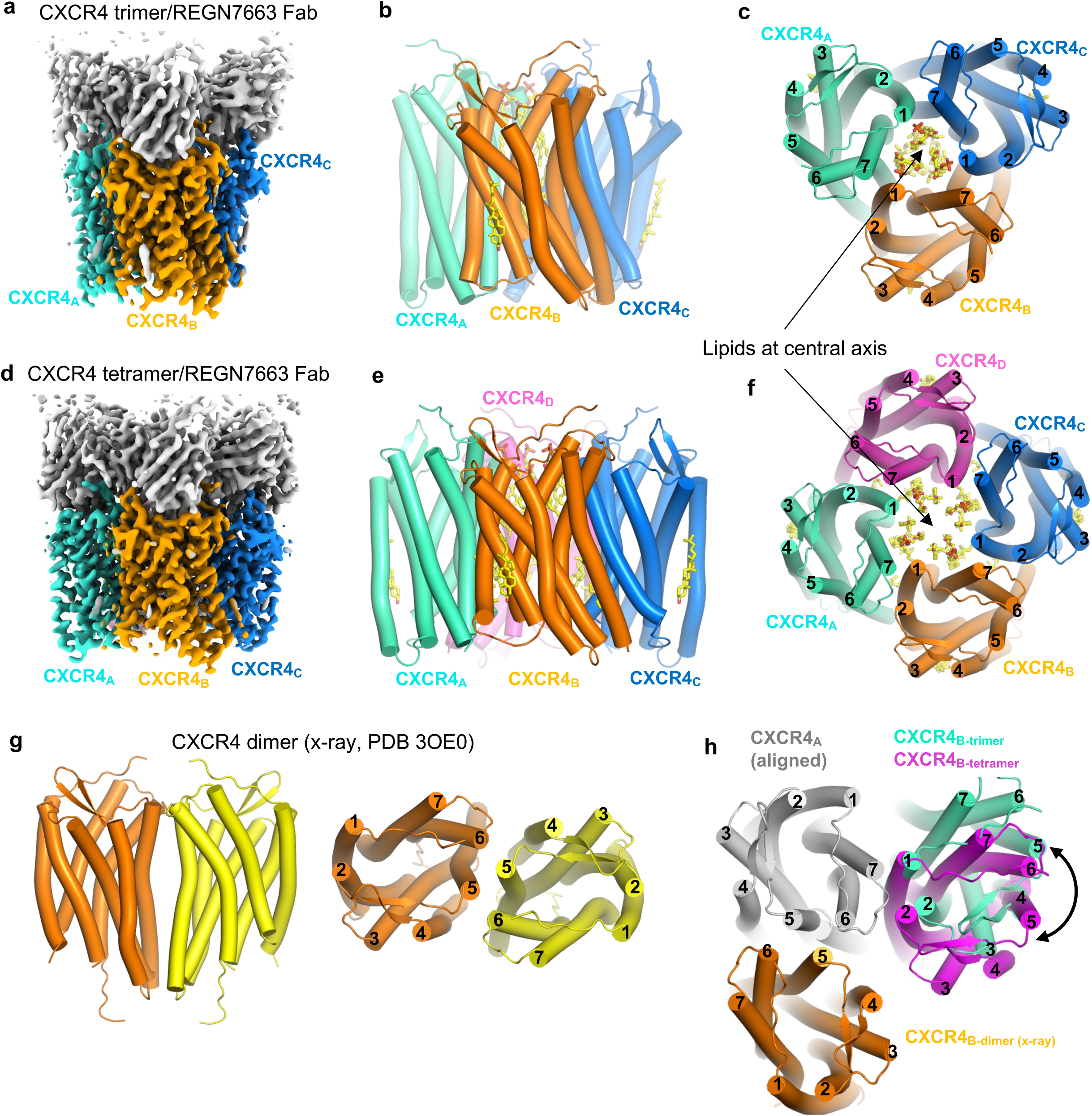
Oligomeric CXCR4 structures. **a**, cryoEM reconstruction of CXCR4 trimer in complex with REGN7663 Fab. **b,c,** side (b) and top-down (c) views of CXCR4 trimer structure. TM helices are shown as cylinders and bound lipids are shown as sticks. Fab molecules are omitted for clarity. **d,** cryoEM reconstruction of CXCR4 tetramer in complex with REGN7663 Fab. **e, f,** side (e) and top-down (f) views of CXCR4 tetramer structure. **g**, side (left) and top (right) views of previously reported dimeric crystal structure of CXCR4. **h**, Top-down view of a CXCR4 protomer (gray) showing positions of neighboring subunits from dimer (orange), trimer (cyan) and tetramer (magenta).

The comparable interprotomer interfaces of trimeric and tetrameric CXCR4 are composed of TM5, TM6, and TM7 of one protomer interacting with TM1 and TM7 of its neighboring protomer (Fig. 5c,f). A ∼20° rotation in the angle between neighboring subunits underlies the distinct oligomeric states (Fig. 5h). This oligomeric interface does not overlap with the dimeric interface observed in crystal structures of CXCR4^19,20^ (Fig. 5h), speculatively allowing for “super-clustering” of CXCR4 protomers mediated by a combination of trimeric/tetrameric and dimeric interfaces (Extended Data Fig. 9a,b). Structural superposition indicates that steric clash caused by the T4L fusion in the crystallization construct may have precluded the assembly of trimers or tetramers observed in our data (Extended Data Fig. 9c,d), thus suggesting why homodimer formation was favored for the T4L-fused receptor.

The trimeric interface is characterized by a buried surface area of ∼1150 Å^2^ and is primarily mediated by crisscrossing of TM6 and TM1 of neighboring protomers near the midpoint of the membrane (Fig. 6a). The diagonal orientation of TM6 results in interprotomer contacts with the cytoplasmic half of TM7. TM1 of the neighboring protomer makes additional interprotomer contacts with cytoplasmic end of TM5 and the extracellular tip of TM7. As expected from interactions between transmembrane helices, most of the residues involved are hydrophobic. As noted above, the tetramer interface is similar to the trimer interface (Fig. 6b). However, close inspection revealed a remarkable difference in the tetramer: a sterol-shaped density that we tentatively built as cholesterol present at the cytoplasmic half of the bilayer sandwiched between TM5/TM6 of one protomer and TM1/TM7 of its neighbor (Fig. 6b,c). To make space for sterol binding at the tetrameric interface, the intracellular portion of TM6 splays away from the interface and TM1 of the neighboring protomer rotates relative to their conformations in the trimeric interface (Fig. 6d). The TM1 rotation is concurrent with the rotation of the entire CXCR4 protomer, which in turn allows space for the additional subunit present in the tetrameric assembly (Fig. 5h). Our structures therefore imply that the absence or presence of lipid at the CXCR4 interprotomer interface may drive the assembly of trimers and tetramers, respectively. These findings provide a structural example supporting the idea that cholesterol regulates chemokine receptor oligomerization^62^.

**Figure 6.**
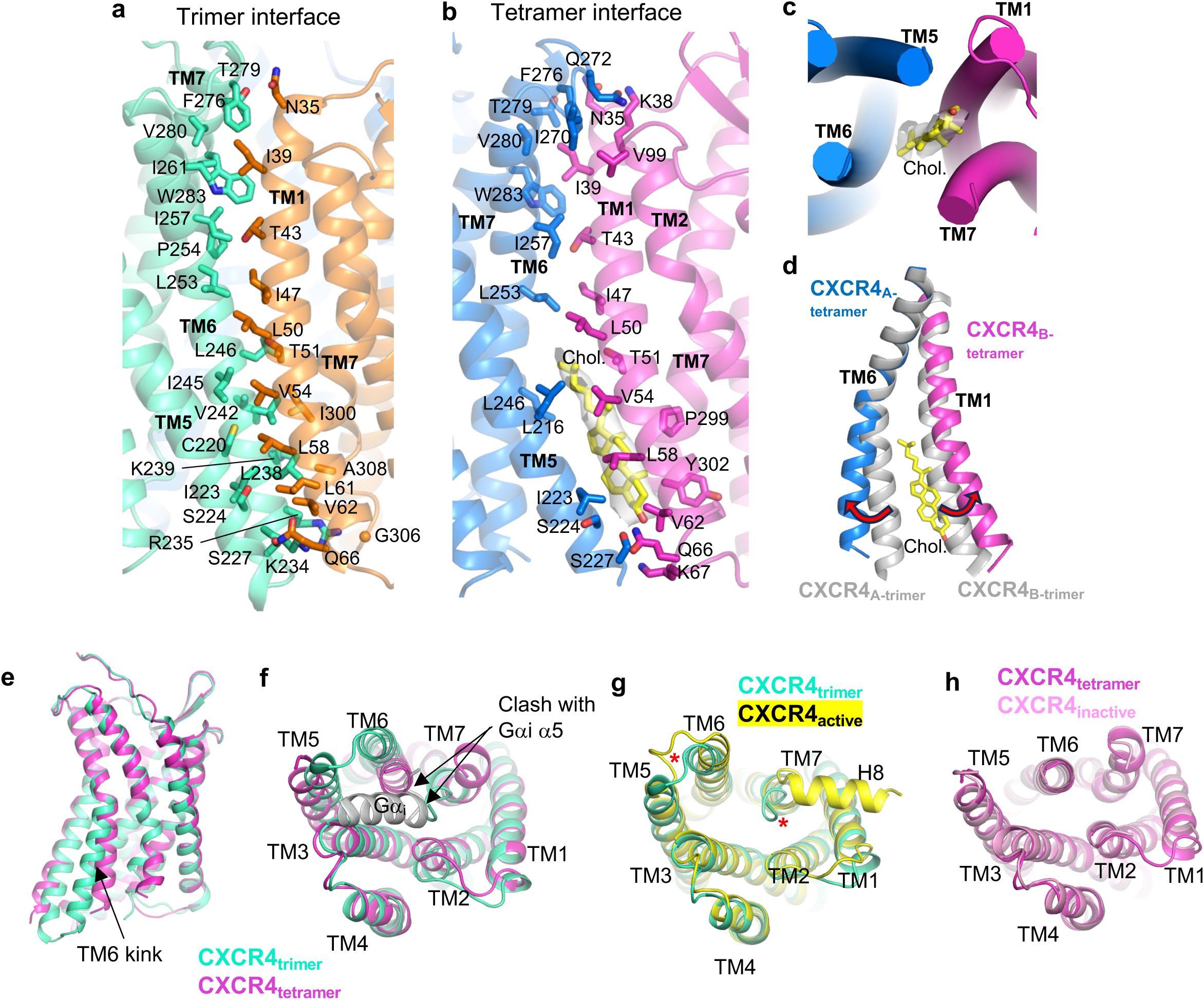
Oligomeric interfaces and protomer conformations. **a**, interprotomer interface of CXCR4 trimer. Interface residues are shown as sticks and labeled. **b,** interprotomer interface of CXCR4 tetramer. Interface residues and modeled cholesterol shown as sticks. Density corresponding to cholesterol is shown as transparent gray surface. **c,** bottom-up view showing position of cholesterol at the tetramer interface. **d,** structural alignment of TM6 and TM1 at the trimer (gray) and tetramer (blue and magenta, with cholesterol in yellow sticks). **e,f,** side (e) and bottom-up (f) views of protomeric structures of trimeric (cyan) and tetrameric (magenta) CXCR4. Binding of Gα_i_ α5 helix (gray) is prevented by steric clash. **g,** structural alignment of trimeric CXCR4 protomer (cyan) and active CXCR4 protomer (yellow). Red asterisks highlight the distinct positions of ICL3 and TM7. **h**, structural alignment of tetrameric CXCR4 protomer (magenta) and inactive CXCR4 (pink).

A super-resolution microscopy study reported that the simultaneous introduction of 3 mutations (K239E/V242A/L246A) within TM6, and located at the oligomerization interface observed in our structures, resulted in reduced higher order oligomerization of CXCR4^21^. We used FSEC to examine the effect of this triple mutant and other mutations at the oligomeric interface on the oligomerization behavior of the detergent-extracted receptor, using CXCR4_EM_ as the background construct (Extended Data Fig. 10a). The K239E/V242A/L246A and K239E/V242W/L246W triple mutants both showed a reduced propensity to form oligomers relative to monomers, determined from the FSEC peak-area ratio of oligomer to monomer for each mutant (Extended Data Fig. 10b). We found that the single mutant V242W showed similarly reduced oligomerization, likely by introducing steric hindrance at the oligomerization interface. On the other hand, L246W increased apparent oligomerization and reduced monomer levels, possibly by augmenting the hydrophobic interactions between subunits. A mutation at a TM1 residue (L58W) that faces TM5 of the neighboring subunit also showed reduced oligomer/monomer ratio. Other TM1, TM6, and TM7 mutants showed no significant change in oligomer/monomer ratio (T51W) or did not have clearly interpretable FSEC chromatograms, presumably due to impacts on expression level or stability of the receptor in detergent. Overall, these biochemical data corroborate the oligomeric interface observed in our structural data.

We next examined the conformations of the individual protomers within the CXCR4 trimer and tetramer. As noted above, a striking difference is the kink at TM6 associated with sterol binding (Fig. 6e,f). TM6 of the trimeric protomer is kinked outward relative to that of the tetrameric protomer, suggesting a more active-like conformation. Indeed, the structure of the trimeric protomer matches closely with active CXCR4 in complex with REGN7663 Fab and G_i_ while the tetrameric protomer aligns well with the inactive CXCR4/REGN7663 Fab complex in the absence of G_i_ (Fig. 6g,h). A noteworthy distinction between the trimeric CXCR4 protomer and active, G_i_-bound CXCR4 is the conformation of ICL3, TM7 and H8; in the trimer, ICL3 is pushed away from the cytoplasmic-facing core, C-terminal end of TM7 is tucked inward, effectively blocking G_i_ binding, and H8 is not visible in the cryoEM map (Fig. 6f,g). Therefore, while trimeric CXCR4 is composed of protomers with an active-like conformation, they are not structurally competent for G_i_-coupling and as such cannot be deemed fully active. This structural observation agrees with FSEC data showing that the presence of G_i_ did not result in a shift of the oligomeric peak (Extended Data Fig. 1b). Overall, these oligomeric structures demonstrate that distinct protomeric conformations underpin the trimeric and tetrameric arrangements of CXCR4. Lipids found at the central axis and at the tetrameric interface appear to be important for oligomeric assembly.

## Discussion

A longstanding drug target for HIV, cancer, and immune disorders, CXCR4 is one of the most well-studied chemokine receptors, and was the first to be crystallized. However, critical structures of CXCR4 remained missing. We have presented here a thorough investigation of CXCR4 structure using cryoEM. Our structure of active CXCR4 bound to CXCL12 shows how the chemokine N-terminus buries deep into the orthosteric pocket to activate the receptor. Mutations at the distal CXCL12 N-terminus^40^ likely diminish its agonistic activity by disrupting the interactions between chemokine and receptor at the TM domain that are required for activation. Due to the flexibility of the complex, we were unable to resolve interactions between the receptor N-terminus and chemokine (chemokine recognition site 1). Therefore, further studies are necessary to visualize this important determinant of CXCL12/CXCR4 affinity.

Like CXCL12, the FDA-approved drug AMD3100 uses electrostatic interactions, namely between its two positively charged lactam rings and acidic residues in the CXCR4 TM domain, to stabilize a diagonal binding mode. We have also shown how a potent antibody inhibitor, REGN7663, blocks CXCL12 by binding across the extracellular face of CXCR4 and partially inserting its CDR-H3 loop into the orthosteric pocket. The structures of REGN7663/CXCR4 complexes do not provide a clear answer as to why this mAb has apparent inverse agonist activity in the setting of CXCR4 overexpression. Stable binding of REGN7663 to active state CXCR4/G_i_, which might be unexpected for an inverse agonist mAb, was possibly enabled by the constitutively active N119S mutation present in our construct that shifts the conformational equilibrium of the receptor. While it is tempting to speculate that inverse agonism is related to interactions between REGN7663 and the CXCR4 TM domains, inverse agonist antibodies raised against the MC4R N-terminus have been reported^63^, suggesting TM domain interactions are not a prerequisite for GPCR inverse agonist mAbs.

Though the functional relevance of chemokine receptor oligomerization *in vivo* awaits confirmation, CXCR4 oligomerization has been reported in various experimental settings, including crystal structures of parallel homodimers^17^. In this study, we have observed that detergent-solubilized CXCR4 exists in various oligomeric states, and determined structures of receptor trimers and tetramers. The parallel orientation of the protomers as well as the encapsulation of lipids at the central axis support the notion that these oligomeric species are present at the cell surface of insect cells overexpressing CXCR4 prior to detergent solubilization. Nonetheless, whether these species correspond to cell membrane oligomers observed previously^16,55^ or are representative of *in vivo* CXCR4 requires further investigation. Interestingly, super-resolution microscopy experiments implicated three TM6 residues (K239, V242, L246) located at the oligomerization interface observed in our structures as being important for higher order oligomerization but not dimerization of CXCR4 in Jurkat cells^21^. Furthermore, the oligomerization-defective K239E, V242A, L246A triple mutant showed decreased chemotaxis in response to CXCL12 *in vitro*^21^. These previously reported data provide a link between our oligomeric structures of detergent-solubilized receptor and CXCR4 function in T cells.

Finally, we observed that oligomeric state and specifically, the binding of lipid at the oligomeric interface, are correlated with distinct conformations of CXCR4 protomers. While the individual protomers of trimeric CXCR4 exhibited an active-like conformation characterized by outward-kinking of TM6, the positioning of intracellular-facing structural elements (ICL3, TM7, and H8) appear to preclude the docking of G_i_. Therefore, additional conformational changes would be required for the oligomeric CXCR4 entities observed here to participate directly in G protein-mediated cellular signaling.

Overall, our structures build on previous crystallographic studies^19,20^ to provide a foundation for understanding how peptides, small molecules, chemokines, and antibody bind and affect the function of CXCR4 in diverse ways. Our data also provide a structural perspective on oligomerization as a potential mode of GPCR regulation, adding a layer of complexity to studies that have focused on monomers as the functional units in physiology and disease.

## Methods

### FSEC-based Construct Screening

Expression constructs (shown in Extended Data Fig. 1a) were codon optimized, synthesized, and cloned into pFastBac1 or pFastbac Dual vectors by Genscript. Second generation baculoviruses (P1) encoding human CXCR4, CXCR4_EM,_ Gα_i_, or Gý_1_/Gψ_2_ (expressed together using pFastBac Dual) were generated in ExpiSf9 cells (ThermoFisher), titered, and adjusted to approximately 2.5×10^8^ ivp/ml. The titering assay was performed using flow cytometry to detect envelope protein gp64 displayed on the surface of infected cells. ExpiSf9 cells at ∼5×10^6^ cells/ml were infected with either CXCR4 alone (1:11 viral dilution), or with Gα_i_ (1:22 viral dilution) and Gý_1_/Gψ_2_ (1:22 viral dilution). Cells were harvested by centrifugation after 72 hr growth (120 rpm shaking, 27°C, 125 ml flat-bottom flask, Innova 44 shaker). After freeze-thaw (−80°C), cell pellets, each from 1 ml of culture, were resuspended in 200 µl lysis buffer (25 mM Tris pH 7.5, 50 mM NaCl, 2 mM MgCl_2_, cOmplete (EDTA-free) protease inhibitor, 5 mM CaCl_2_, 50 mU/mL Apyrase) and rotated at 4°C for 1 hr. For the samples to which G_i_ was added, G_i_ containing pellets were first suspended in 200 µl lysis buffer. 200 µl of G_i_ slurry was then used to resuspend the receptor containing pellets. After 1 hr, 200 µl of solubilization buffer (25 mM Tris pH 7.5, 50 mM NaCl, 2 mM MgCl_2_, 5 mM CaCl_2_, ∼2% LMNG, ∼0.2% CHS, cOmplete (EDTA-free) protease inhibitor, 50 mU/ml Apyrase) was added and the mixture was rotated at 4°C for an additional 1 hr at 4°C. Insoluble material was removed by centrifugation and each sample was subjected to FSEC (buffer: 25 mM Tris pH 7.5, 150 mM NaCl, 2 mM MgCl2, 0.01% LMNG, 0.001% CHS). A Zenix-C SEC-300 3 µM 300 Å 4.6×300mm column (flow rate: 0.35 ml/min) was used for the data shown in Extended Data Fig. 1b. For the data shown in Extended Data Fig. 10, a Zenix-C SEC-300 3 µM 300 Å 7.8×300mm column (flow rate: 0.75 ml/min) was used and the baculovirus used was not titered. FSEC data were collected using a Shimadzu LC system using LabSolutions v5.111 software.

### Expression and Purification of CXCR4 and G_i_ proteins

ExpiSf9 cells at ∼5×10^6^ cells/ml were infected with P1 baculovirus encoding either CXCR4_EM_ or Gα_I_ and Gý_1_/Gψ_2_ as described above. Cells were harvested by centrifugation (3000 x g, 10 min, 4°C) after 72 hr growth (120 rpm shaking, 27°C, 2 L flat-bottom flask, Innova 44 shaker). Cell pellets were washed in ice-cold DPBS with cOmplete (EDTA-free) protease inhibitor and then subjected to freeze-thaw (−80°C) then resuspended in lysis buffer (25 mM Tris pH 7.5, 50 mM NaCl, 2 mM MgCl_2_. 1x cOmplete (EDTA-free) protease inhibitor, 5 mM CaCl_2_, 50 mU/mL Apyrase). Crude lysates containing CXCR4_EM_ and G_i_ were then combined and stirred at 4°C. After 1 hr, an equal volume (1 ml for every 1 ml of lysis buffer) of solubilization buffer (25 mM Tris pH 7.5, 50 mM NaCl, 2 mM MgCl_2_, 5 mM CaCl_2_ 2% LMNG, 0.2% CHS) was added to the slurry and the mixture was stirred at 4°C for 1 hr. Insoluble material was removed by centrifugation (100,000 x g, 4°C, 30 min). Anti-FLAG M2 Affinity Gel (Sigma cat# A2220) was used to capture CXCR4_EM_-containing species. The protein-loaded resin was washed with SEC buffer (25 mM Tris pH 7.5, 150 mM NaCl, 2 mM MgCl_2_, 0.01% LMNG, 0.001% CHS) and protein was eluted in SEC buffer containing 0.15 mg/ml 3x FLAG peptide. The eluate was concentrated to approximately 0.5 ml and subjected to SEC. A tandem column was used to improve separation of different CXCR4_EM_ species: a Superose 6 Increase 10/300 GL column was connected upstream of a Superdex 200 Increase 10/300 GL column. Fractions containing CXCR4_EM_/G_i_ protein complex were selected, pooled, concentrated, and mixed with either Fab’, CXCL12, or AMD3100 prior to cryoEM grid making.

A comparable procedure was used for the production CXCR4_EM_ to which G_i_ was not added. In this case, SEC peaks corresponding to oligomeric and monomeric CXCR4_EM_ were separately harvested and were each mixed with Fab’ prior to cryoEM grid making.

### Fab’ Production

REGN7663 IgG was diluted to 2 mg/ml in 20 mM HEPES pH 7.4, 150 mM NaCl. IdeS, an IgG-specific protease, was added to cleave off Fc region thereby producing F(ab’)_2_. 10 µg concentrated IdeS per 1 mg antibody (1:100) was added and the cleavage reaction was carried out at 37°C for 30 min. F(ab’)_2_ was reduced using approximately 88 mM cysteamine hydrochloride at 37°C for 10 min, in the presence of approximately 18 mM EDTA. Reduced Fab’ was dialyzed against 20 mM HEPES pH 7.4, 150 mM NaCl overnight at 4°C. Fab’ was further purified by IMAC (negative-pass to remove His-tagged IdeS) and CaptureSelect IgG-Fc (Multispecies) Affinity Matrix (negative-pass to remove Fc fragment.) F(ab’) was treated with 20 mM iodoacetamide at room temperature, in the dark, for 30 min to alkylate the reduced hinge cysteines. Fab’ was purified further via SEC (HighLoad 16/600 Superdex 75 pg column equilibrated to 25 mM Tris pH 7.5, 150 mM NaCl), and concentrated before use.

### CRE-Luciferase CXCR4 functional assay

HEK293 cell lines were generated to stably express full-length human CXCR4 (hCXCR4; amino acids 1-352 of accession number NP_003458.1) along with a luciferase reporter cAMP response element (CRE, 4X)-luciferase-IRES-GFP. For CXCR4 CRE-Luciferase assay, HEK293/CRE-Luc/hCXCR4 cells were plated in Opti-MEM media (Invitrogen, cat# 31985-070) supplemented with 0.1% FBS (Seradigm, Cat#1500-500) at 37°C with 5% CO_2_ for overnight. The cells were then incubated with 5uM of Forskolin (Sigma, cat# F6886) and serially diluted CXCL12 (Tocris, Cat# 350-NS) for activation of CXCR4 or pre-incubated with REGN7663 or control antibody for 30 minutes before adding 5uM of Forskolin without or with 500pM of SDF for inhibition of CXCR4 basal activity or SDF-induced CXCR4 activation. Cells were incubated for 5.5 hours at 37°C with 5% CO_2._ At the conclusion of the incubations, the luciferase activity was detected using OneGlo (Promega, Cat# E6130) and luminescence was recorded by an EnVision Plate reader using EnVision Manager v1.14 (Perkin Elmer). Results were analyzed using nonlinear regression (4-parameter logistics) with Prism 6 software (GraphPad) to obtain EC_50_ and IC_50_ values.

### CryoEM grid preparation and data collection

CXCR4_EM_ (G_i_-bound complex, monomer, or oligomer) were concentrated to ∼1 to ∼5 mg/mL and left as is (“apo”, G_i_-bound complex sample) or mixed with 0.5 mg/mL CXCL12 (Recombinant Human/Rhesus Macaque/Feline CXCL12/SDF-1 alpha, R&D Systems Catalog #: 350-NS-050/CF), or 1 mM AMD3100 (AMD 3100 octahydrochloride, R&D Systems Catalog #: 3299), or ∼1 to 1.5 mg/mL REGN7663 Fab and incubated on ice for ∼1 hour. Samples were pipetted onto freshly hydrogen/oxygen plasma cleaned UltrAuFoil 0.6/1 300 mesh grids and blotted then plunge frozen into liquid ethane using a Vitrobot Mark IV and stored in liquid nitrogen prior to data collection.

Samples were inserted into a Titan Krios G3i (ThermoFisher) microscope equipped with a BioQuantum K3 (Gatan) imaging system or a Glacios microscope equipped with a Falcon 4i camera and Selectris energy filter (ThermoFisher). Data were collected at nominal magnifications of 105 kx (0.85 Å/pixel) or 165 kx (0.696 Å/pixel) and energy filters were inserted with slit widths of 20 ev and 10 ev on the Titan Krios and Glacios microscopes, respectively. Automated data collections were carried using EPU v2.12 with an applied defocus range of -1.0 to -2.2 µM. A 40° stage tilt was applied during collection of the oligomeric CXCR4_EM_/REGN7663 Fab complex sample to overcome preferred particle orientations. Additional details regarding data collection are shown in Extended Data Table 1.

### CryoEM image processing

CryoEM data processing for apo CXCR4_EM_/G_i_, CXCR4_EM_/G_i_/AMD3100 CXCR4_EM_/G_i_/REGN7663Fab, CXCR4_EM_/REGN7663 Fab trimer, and CXCR4_EM_/REGN7663 Fab tetramer was carried out within the cryoSPARC v3.3.2 pipeline^64^. Patch motion correction and Patch CTF estimation were used to align movie frames and estimate CTF parameters, respectively. Particle images were picked using 2D template based picker or TOPAZ v0.2.5^65^ then extracted and subjected to multiple rounds of 2D classification, ab initio reconstruction and heterogeneous refinement to obtain a homogenous subset of particles with well resolved features corresponding to the target complex. Final map calculations were carried out using the Local Refinement job type. C3 and C4 symmetry were applied for refinement of the trimeric and tetrameric reconstructions of CXCR4_EM_/REGN7663 Fab, respectively. Refinements of oligomeric CXCR4 conducted without applied symmetry yielded similar structures to the symmetric refinements, but at lower resolution.

Initial processing steps for the CXCR4_EM_/G_i_/CXCL12 and CXCR4_EM_/REGN7663 Fab monomeric complexes were carried out in RELION-3^66^. CTF parameters were calculated using gctf^67^ and CTFFIND4^68^. Particles were picked using TOPAZ^65^, then sorted by 2D and 3D classification. Initial 3D refinements of the CXCR4_EM_/G_i_/CXCL12 complex had very weak density for the ligand. To improve signal for the bound ligand, successive rounds of alignment-free focused 3D classification was conducted, applying a mask around CXCL12. Selected particle images were then subjected to Bayesian polishing and then imported into cryoSPARC for final map refinements. For the CXCR4_EM_/REGN7663 Fab complex, signal from constant region of the Fab was subtracted prior to final local refinement in cryoSPARC. Additional data processing details are listed in Extended Data Table 1.

### Model building, structure refinement, and visualization

Model building was initiated by docking starting models into the cryoEM maps using the fit in map function in Chimera^69^, followed by rounds of manual adjustment in coot 0.8.9^70^ and real space refinement in Phenix 1.19^71^. Published structures of CXCR4 (PDB 4RWS^20^), G_i_ heterotrimer (PDB 7T2G), and an internal Fab structure were used as initial models to build the CXCR4_EM_/G_i_/REGN7663 Fab complex. CXCR4_EM_ and G_i_ from this structure was then used as starting models for the other structures in this study. A crystal structure of CXCL12 (PDB 3HP3^72^) was used as an initial model for the chemokine. Side chains for CXCL12 residues 13-65 (excepting disulfide bonds) were truncated to C_ý_ due to weak density. The REGN7663 Fab constant regions were omitted from the CXCR4_EM_/REGN7663 Fab (without G_i_), CXCR4_EM_/REGN7663 Fab trimer, and CXCR4_EM_/REGN7663 Fab tetramer models due to weak density. The eLBOW program^73^ in Phenix was used to generate ligand coordinates and restraints for AMD3100. Structures were validated using Phenix and MolProbity^74^. Buried surface areas were calculated using PISA^75^. Pymol^76^, Chimera version1.16^69^, and ChimeraX version 1.2.5^77^ were used to visualize structural data and generate figures.

## Data and materials availability

Regeneron materials described in this manuscript may be made available to qualified, academic, noncommercial researchers through a materials transfer agreement upon request at https://regeneron.envisionpharma.com/vt_regeneron/. For questions about how Regeneron shares materials, use the email address. Atomic coordinates and cryoEM maps have been deposited into the Protein Data Bank (PDB) and Electron Microscopy Data Bank (EMDB) under the respective accession codes 8U4N and 41888 (Apo CXCR4_EM_/G_i_), 8U4O and 41889 (CXCR4_EM_/G_i_/CXCL12), 8U4P and 41890 (CXCR4_EM_/G_i_/AMD3100), 8U4Q and 41891 (CXCR4_EM_/G_i_/REGN7663Fab), 8U4R and 41892 (CXCR4_EM_/REGN7663 Fab), 8U4S and 41893 (CXCR4_EM_/REGN7663 Fab Trimer), 8U4T and 41894 (CXCR4_EM_/REGN7663 Fab Tetramer).

## Acknowledgements

We thank various Regeneron scientists including Yi Zhou, Micah Rapp, and Drew Murphy for discussions, Linda Molla and Samira Chandwani for project management, and Regeneron cloud/HPC teams for supporting cryoEM data storage and processing.

## Author contributions

K.S., L.L.M., J.H., M.M., J.H.K. and M.C.F. conceptualized the studies. L.L.M. and T.R. expressed and purified proteins for cryoEM. K.S. conducted cryoEM experiments and analyzed structural data, with contributions from M.C.F.. J.H. and S.S. conducted Cre-Luciferase assays. K.S., L.L.M., J.H., M.M., J.H.K., R.L., W.C.O., and M.C.F. analyzed data and supervised the overall project. K.S. and L.L.M. drafted the manuscript with input from J.H. and M.C.F. The manuscript was finalized by all authors.

## Competing interests

Regeneron authors own options and/or stock of the company. W.C.O. is an officer of Regeneron.

---

**Extended Data Fig. 1.**
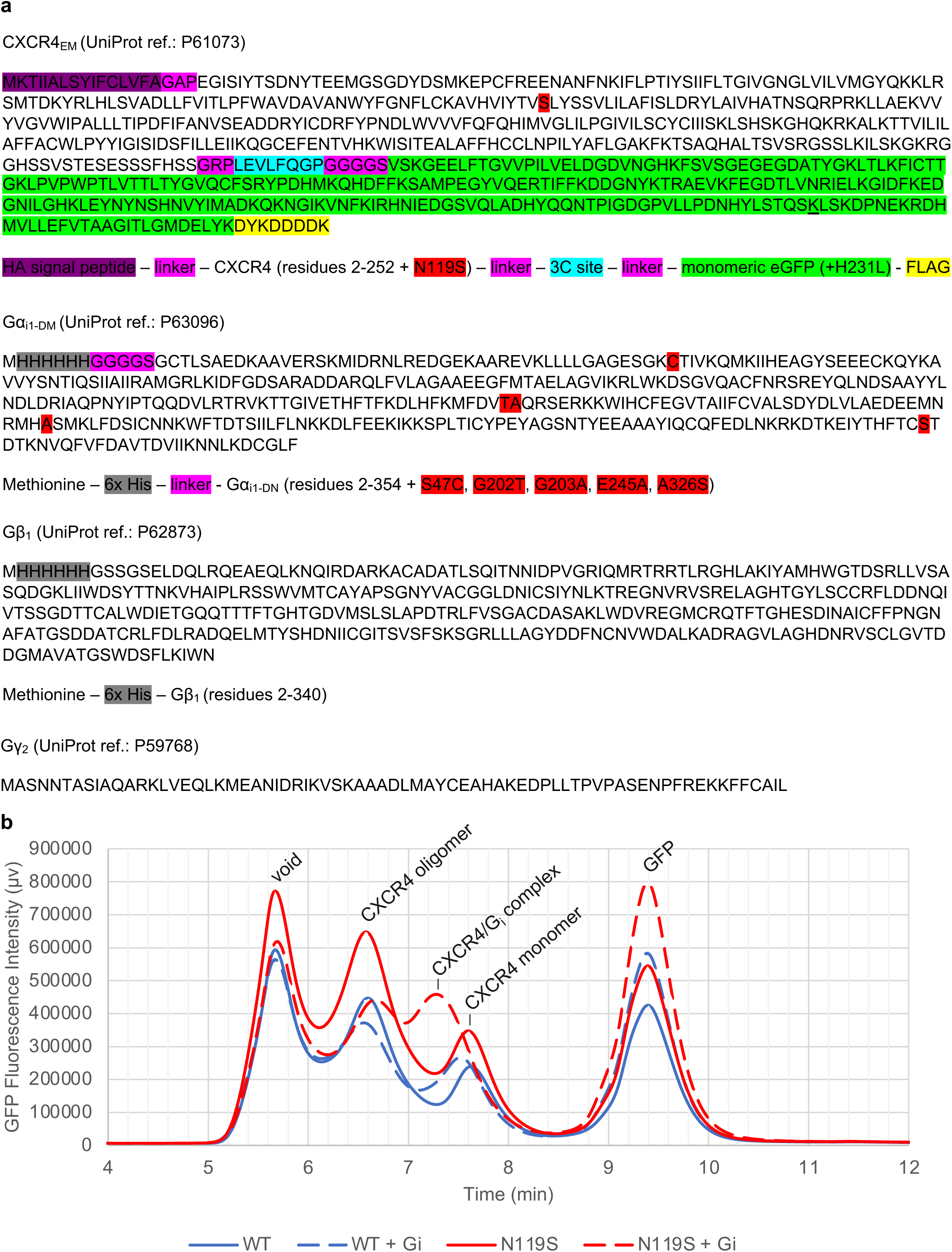
Protein constructs and FSEC-based protein screening. **a**, Primary structures of protein constructs used in structural studies. **b,** Fluorescence-detection size exclusion chromatography (FSEC) screening of wild type CXCR4 (blue) and N119S-containing CXCR4 (referred to as CXCR4_EM_, red) in the presence (solid lines) or absence (dashed lines) of added G_i_. The nominally wild type CXCR4 construct was identical to that shown in (a) without the N119S point mutation. Chromatograms are annotated with presumed peak positions of various species present in the samples.

**Extended Data Fig. 2.**
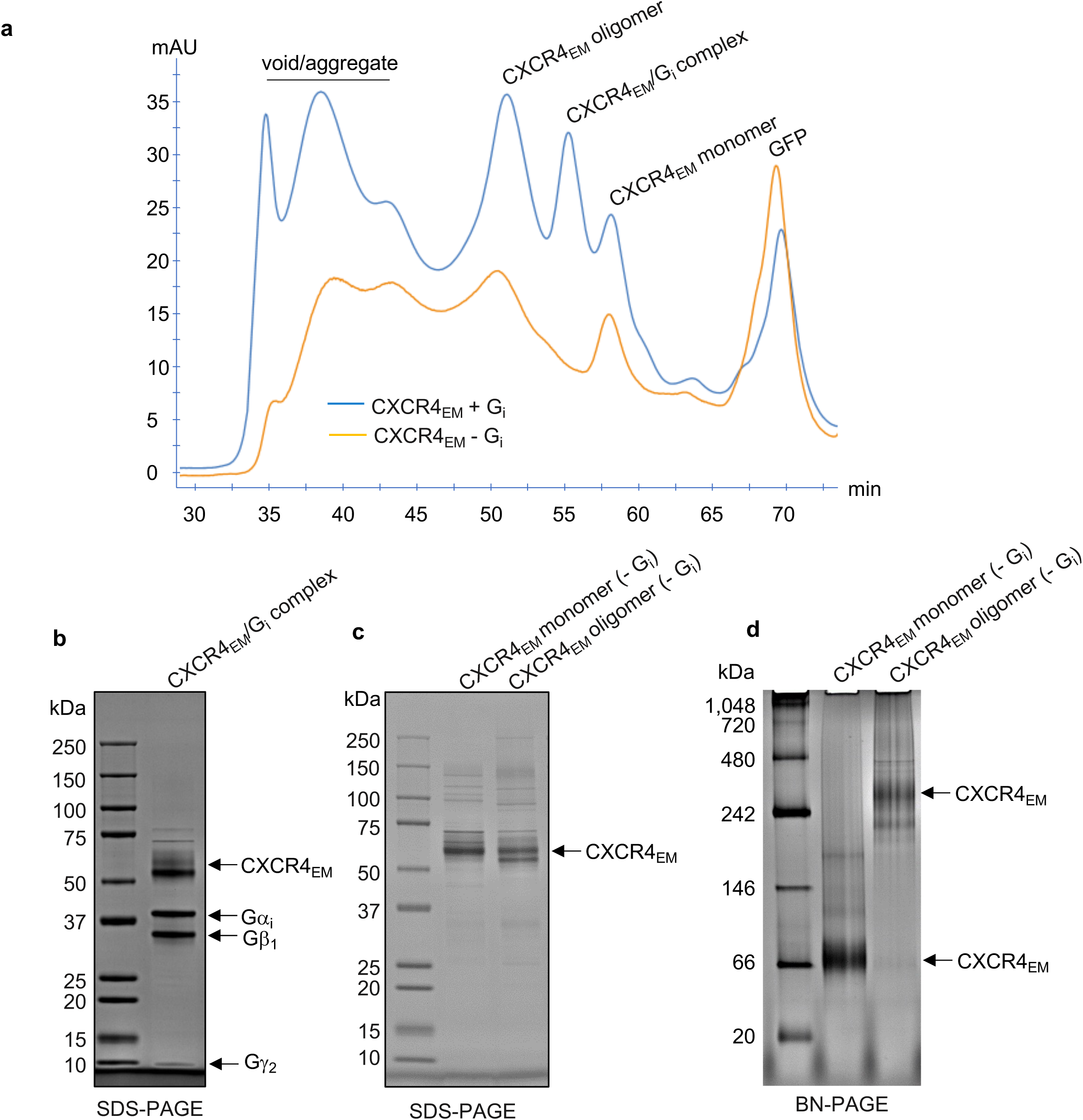
Purification of CXCR4 complexes. **a**, Size-exclusion chromatography (SEC) traces for a CXCR4_EM_ purification with added G_i_ (blue), and for a purification prepared in the absence of exogenously added G_i_ (orange). **b,c** SDS-PAGE (4-20% Tris-Glycine) showing the subunit content and purity of prepared cryoEM samples for CXCR4_EM_/G_i_ complex (b) and CXCR4_EM_ prepared in the absence of added G_i_. CXCR4_EM_/G_i_ complex sample is representative of multiple purifications performed. 2% (v/v) 2-Mercaptoetanol was present in the SDS-PAGE samples prior to loading. **d,** Blue native (BN) PAGE (4-16% Bis-Tris) of SEC-purified CXCR4_EM_ monomer and oligomer samples prepared in the absence of added G_i_.

**Extended Data Fig. 3.**
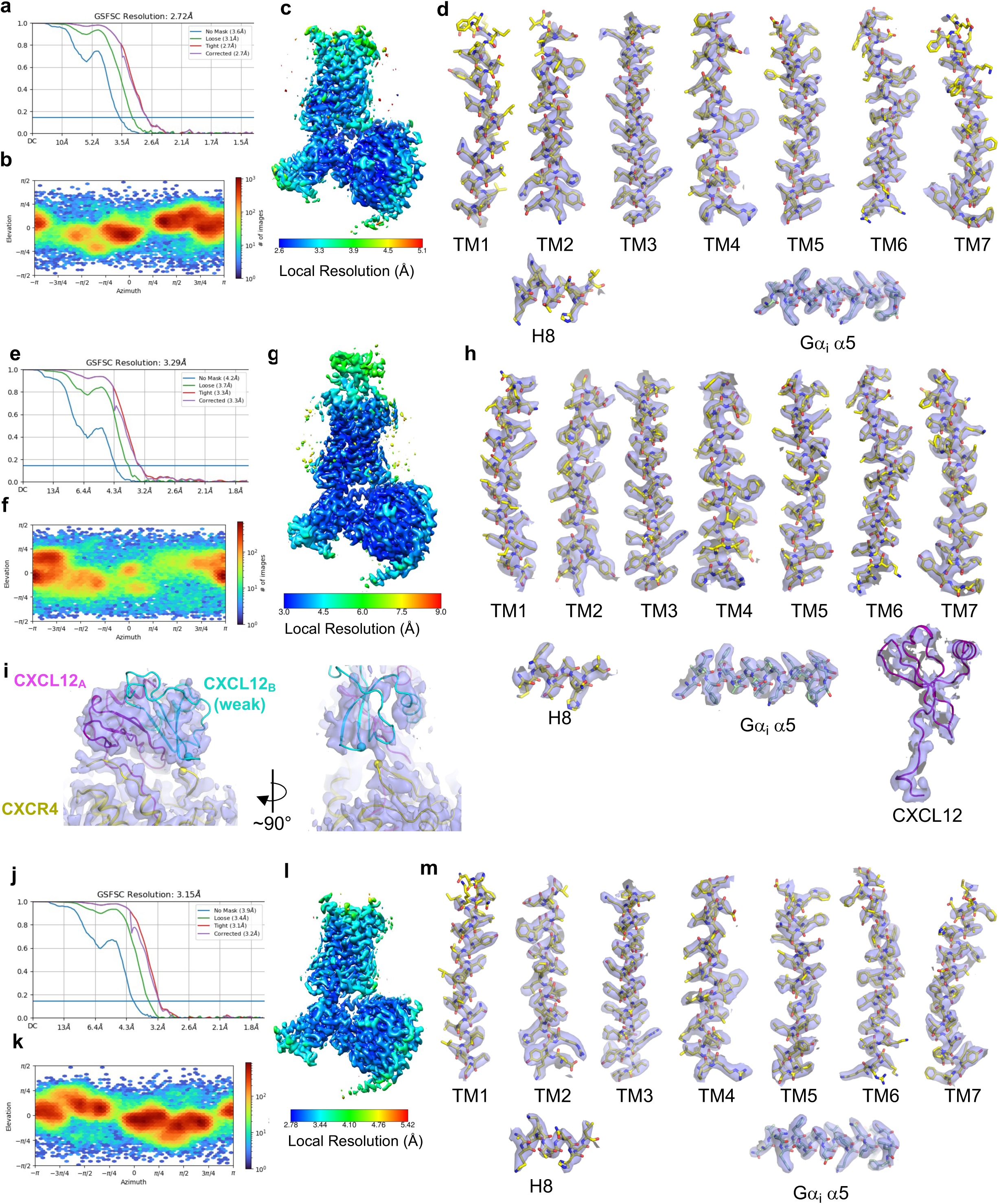
CryoEM reconstruction of Apo CXCR4_EM_/G_i_, CXCL12-bound CXCR4_EM_/G_i_, AMD3100-bound CXCR4_EM_/G_i._ **a-d**, FSC curve (a), particle angular distribution plot (b), local resolution map calculated in cryoSPARC (c), and map/model fits of selected regions (d) for Apo CXCR4_EM_/G_i_. **e-h**, FSC curve (e), particle angular distribution plot (f), local resolution map calculated in cryoSPARC (g), and map/model fits of selected regions (h) for CXCL12-bound CXCR4_EM_/G_i_. **i**, two views showing fit of a CXCL12 dimer (arranged on the basis of PDB 3GV3) into cryoEM map, shown at 4 sigma in pymol. **j-m,** FSC curve (j), particle angular distribution plot (k), local resolution map calculated in cryoSPARC (l), and map/model fits of selected regions (m) for AMD3100-bound CXCR4_EM_/G_i_.

**Extended Data Fig. 4.**
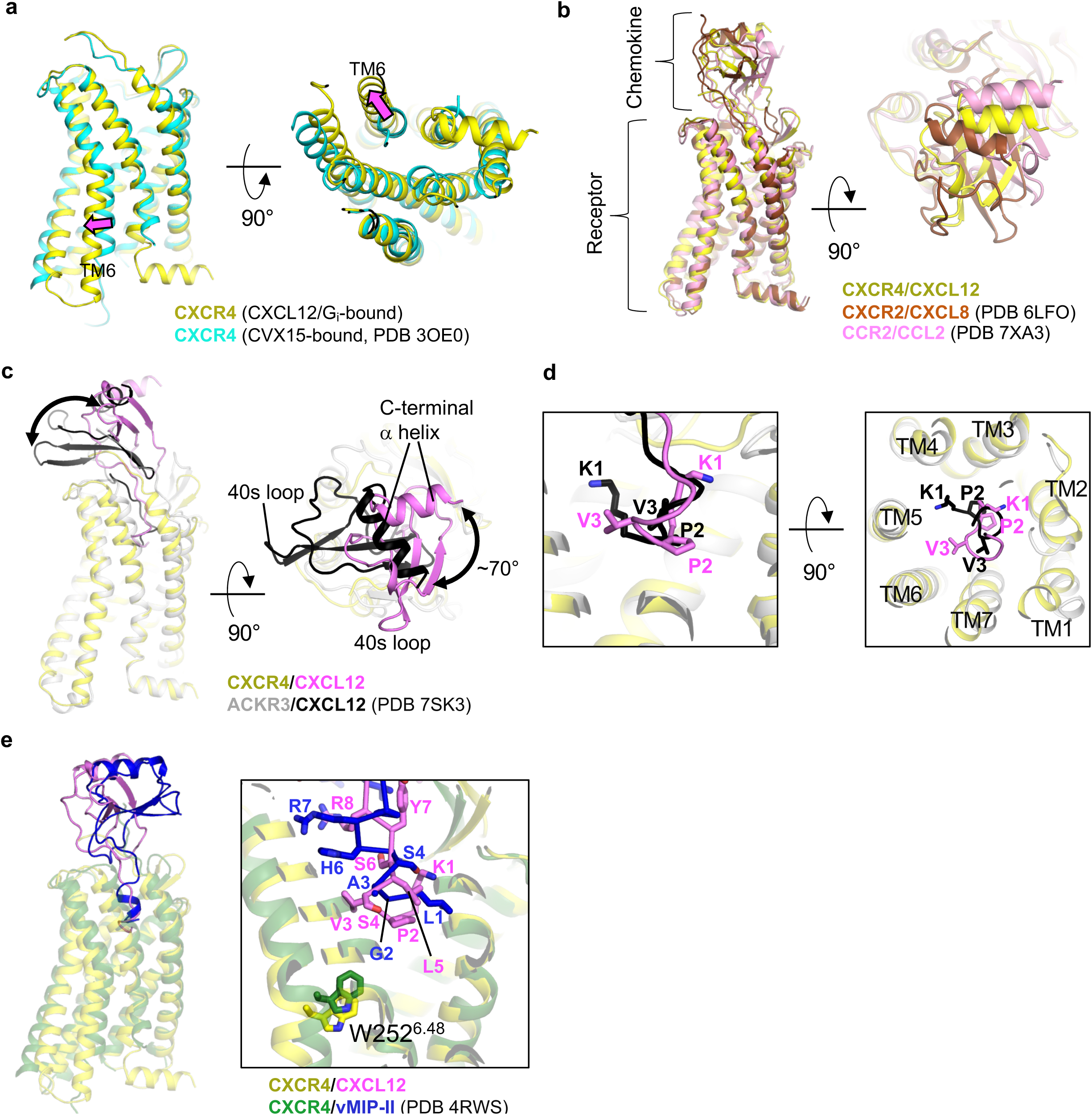
Structural comparisons of chemokine receptor structures. **a**, structural alignment of active CXCR4 (yellow, this study, CXCL12/G_i_-bound complex) and inactive, antagonist-bound CXCR4 (cyan, PDB 3OE0). Magenta block arrows depict movement of TM6. **b**, Alignment of CXCR4/CXCL12 complex (yellow) with CXCR2/CXCL8 complex (brown, PDB 6LFO), and CCR2/CCL2 complex (pink, PDN 7XA3). G protein models are omitted for clarity. **c**, Receptor-based alignment of CXCR4/CXCL12 (yellow/pink) with ACKR3/CXCL12 (gray/black, PDB 7SK3). Arrow highlights different docking orientations of CXCL12 onto the two receptors. **d**, expanded views of showing different binding modes of CXCL12 N-termini (pink in CXCR4 complex and black in ACKR3 complex) in CXCR4 and ACKR3. **e**, alignment of CXCR4/CXCL12 complex (yellow/pink) and CXCR4/vMIP-II (green/blue). Inset shows expanded views of chemokine N-terminal positions within orthosteric pocket, highlighting the positions of toggle switch residue W252 in sticks.

**Extended Data Fig. 5.**
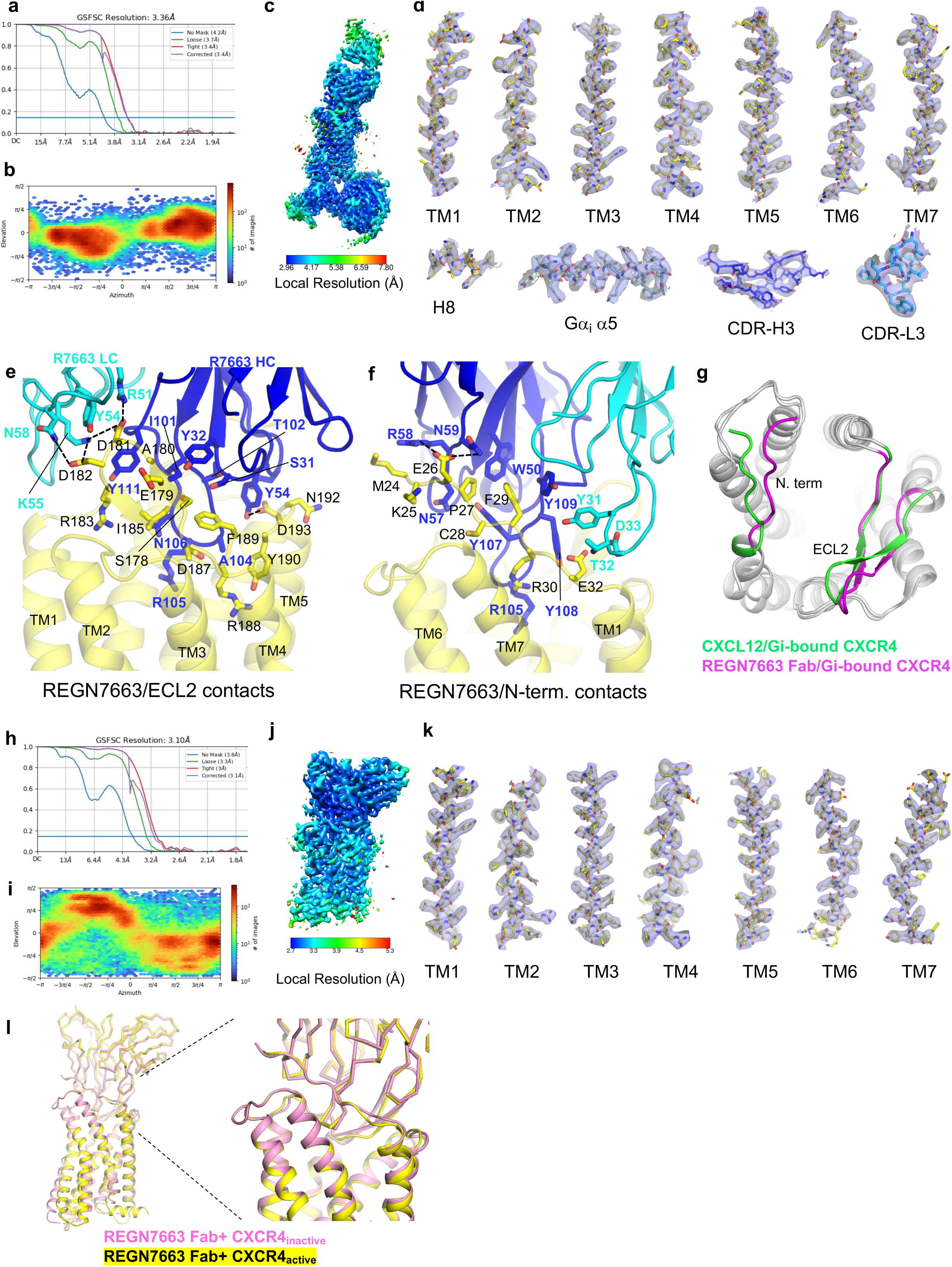
CryoEM reconstruction of REGN7663 Fab/CXCR4_EM_/G_i_ and REGN7663 Fab/CXCR4_EM._ **a-d**, FSC curve (a), particle angular distribution plot (b), local resolution map calculated in cryoSPARC (c), and map/model fits of selected regions (d) for REGN7663 Fab/CXCR4_EM_/G_i_. **e,f,** Expanded view of contacts between REGN7663 Fab (light chain in cyan, heavy chain in blue) and CXCR4 ECL2 (e) and N-term (f). Epitope and paratope residues are shown as sticks and labeled, and apparent salt bridges/hydrogen bonds between mAb and receptor are shown as dashed lines. **g**, structural alignment of CXCR4 bound to CXCL12 and REGN7663 Fab. N-term. and ECL2 are colored green (CXCL12-bound) or magenta (REGN7663 Fab-bound) to highlight their different positions. **h-k**, FSC curve (h), particle angular distribution plot (i), local resolution map calculated in cryoSPARC (j), and map/model fits of TM helices (k) for REGN7663 Fab/CXCR4_EM_ without G_i_. **i**, aligned structures of CXCR4/REGN7663 Fab complex in the inactive (pink) and active (yellow, Gi-bound) conformations. Note the REGN7663 Fab variable region and cytoplasmic half of the CXCR4 domain are mostly superimposable.

**Extended Data Fig. 6.**
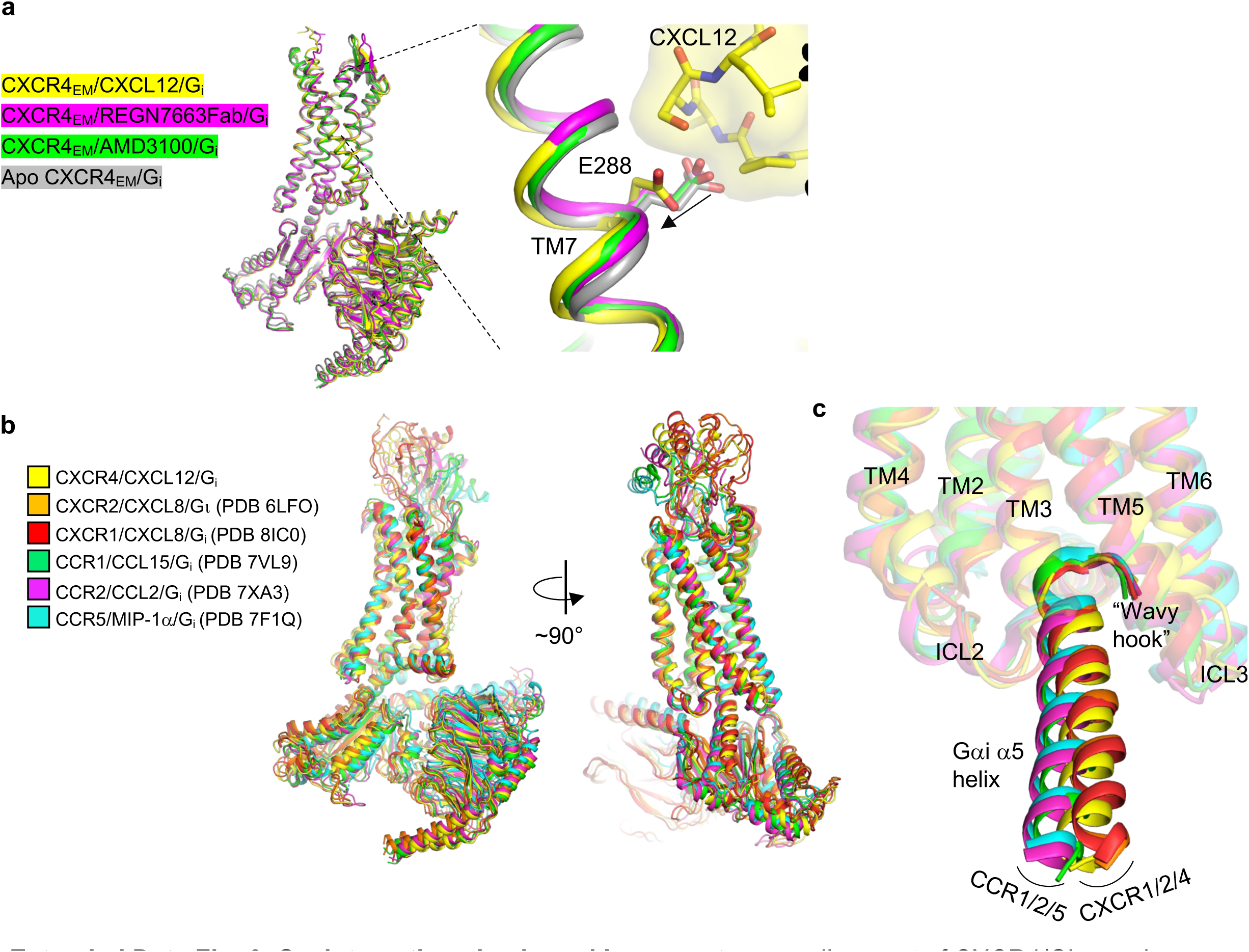
Gα_i_ interactions in chemokine receptors. **a**, alignment of CXCR4/Gi complexes bound to CXCL12 (yellow), REGN7663 Fab (magenta), AMD3100 (green) or in the absence of ligand (apo, gray). Inset shows expanded view around E288 residue. Bound CXCL12 is shown as yellow transparent surface and sticks, highlighting how it enforces a rotameric change of E288 and slight shift of TM7 in the CXCL12-bound complex. **b,** receptor-based alignment showing architecture of various chemokine/chemokine receptor/Gi complexes. **c,** expanded view showing docking of Gαi α5 helix into cytoplasmic pocket of chemokine receptors. Note that the Gαi α5 helix is positioned closer to ICL2 in CC chemokine receptor complexes, while it is closer to ICL3 in CXC chemokine receptor complexes.

**Extended Data Fig. 7.**
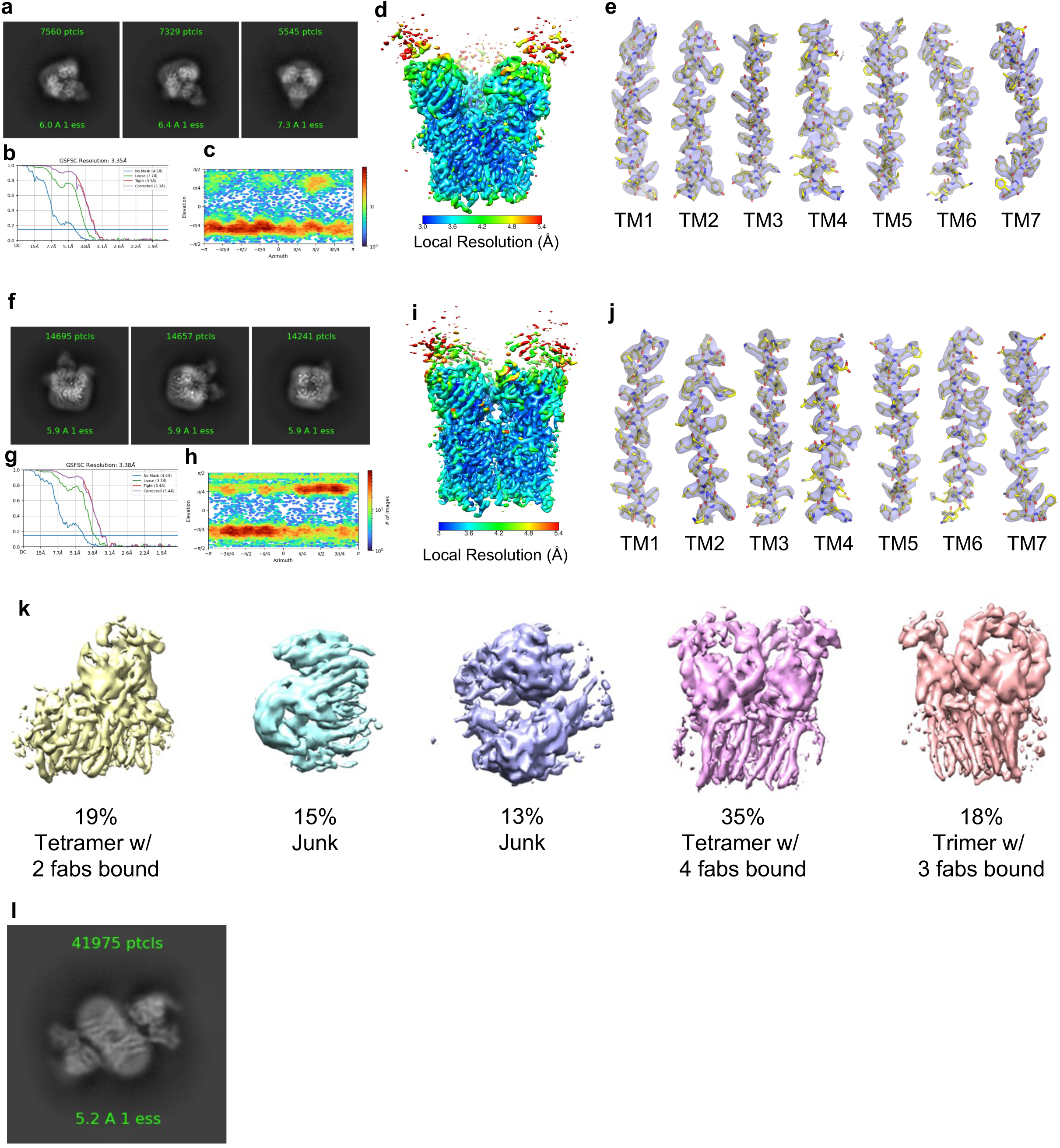
CryoEM of CXCR4 oligomers. **a-e**, example 2D class averages obtained from tilted data collection (a), FSC curves (b), angular distribution plot (c), local resolution map (d) and model/map fits of TM helices (e) of trimeric CXCR4_EM_/REGN7663 Fab complex. **f-j,** example 2D class averages obtained from tilted data collection (f), FSC curves (g), angular distribution plot (h), local resolution map (i) and model/map fits of TM helices (j) of tetrameric CXCR4_EM_/REGN7663 Fab complex. **k**, Output maps from ab initio reconstruction conducted on the oligomeric CXCR4_EM_/REGN7663 Fab particles. Particles belonging to classes of tetramer with 4 fabs bound or trimer with 3 fabs bound were selected for further processing. **l**, An example 2D class average showing an anti-parallel dimer of CXCR4_EM_/G_i_.

**Extended Data Fig. 8.**
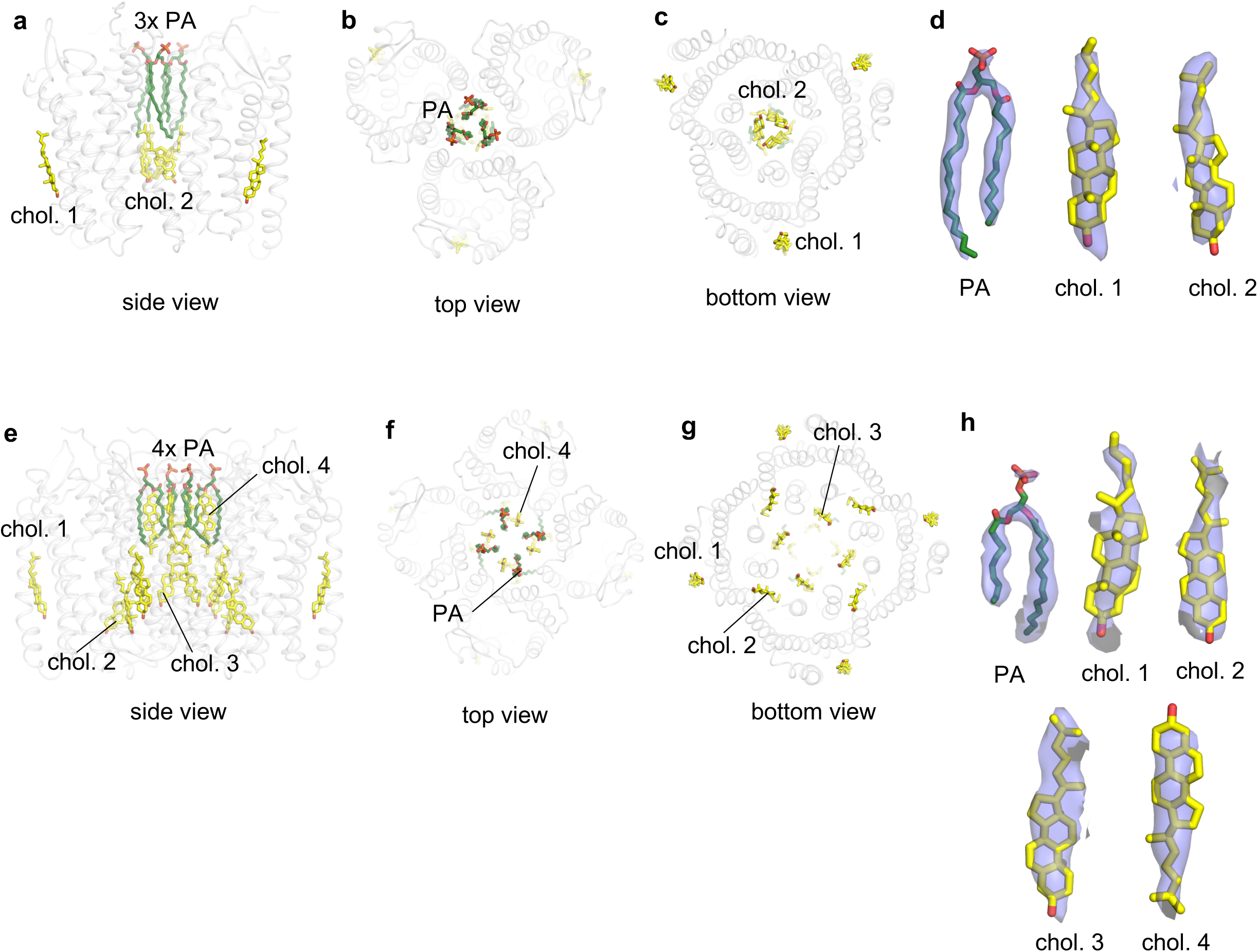
Lipids resolved in oligomeric structures of CXCR4. **a-c**, side (a), top-down (b) and bottom-up (c) views of trimeric CXCR4, highlighting positions of built lipid molecules. Cholesterols (chol.) are shown as yellow sticks and phosphatidic acid (PA) are shown as green sticks. **d,** fit of lipid molecules (shown as sticks) to map (transparent blue surface) in trimeric CXCR4. Chol. 1 and chol 2. refer to lipids labeled in a-c**. e-g,** side (e), top-down (f) and bottom-up (g) views of tetrameric CXCR4, highlighting positions of built lipid molecules. **h,** fit of lipid molecules (shown as sticks) to map (transparent blue surface) in tetrameric CXCR4.

**Extended Data Fig. 9.**
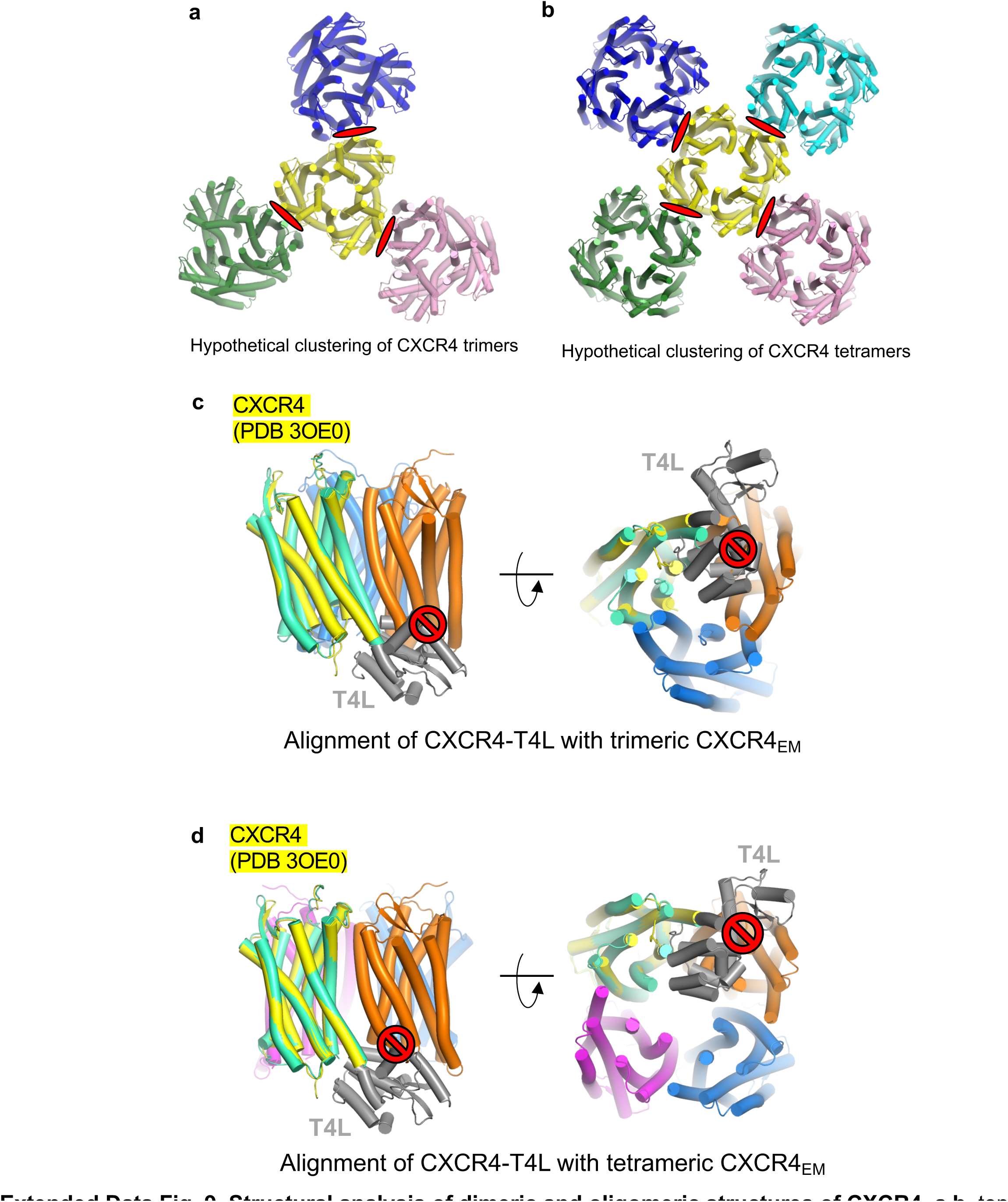
Structural analysis of dimeric and oligomeric structures of CXCR4. **a,b**, top-down view of hypothetical models of four CXCR4 trimers (a) or five CXCR4 tetramers (b) clustered via dimeric interfaces (red ovals) observed in crystal structures. **c,d,** a single subunit from dimeric x-ray structure of CXCR4 (receptor in yellow, ICL3-fused T4L in gray) aligned to trimeric CXCR4 (c) or tetrameric CXCR4 (d). Side and top views are shown. Red symbols indicate steric clash between T4L and neighboring protomers that would prevent trimer or tetramer assembly. The steric hindrance caused by fused T4L may explain why dimeric CXCR4 was favored over trimeric or tetrameric CXCR4 in previous crystallographic studies.

**Extended Data Fig. 10.**
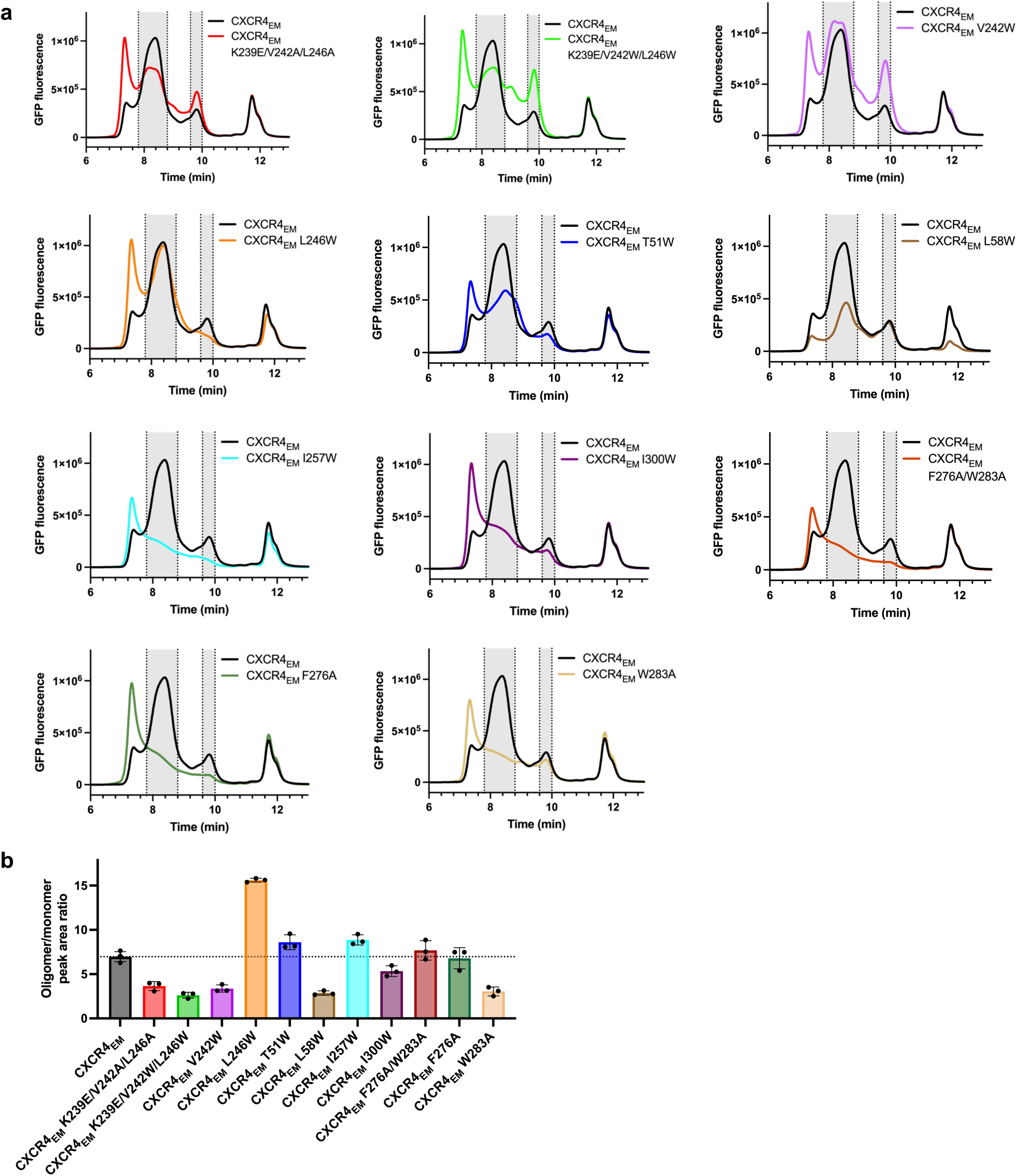
FSEC analysis of CXCR4 oligomerization interface mutants. **a**, FSEC chromatograms of control construct CXCR4_EM_ (black trace) and its mutants (colored traces) tracking GFP fluorescence. The same chromatogram for the control are shown in each for comparison. Gray shaded regions indicate elution times corresponding to CXCR4 oligomer (7.8-8.8 min) and monomer (9.6-10 min). **b**, ratio of oligomer to monomer peak areas, calculated according to shaded regions in **a**. horizontal dotted line corresponds to mean value for CXCR4_EM_. Column heights indicate mean values, and error bars show standard deviations calculated from N=3 or 4 FSEC experiments using two independently generated baculoviruses for each construct. Note that several mutants (I257W, I300W, F276A/W283A, F276A, W283A) showed poor chromatographic behavior overall, presumably due to poor expression or stability in detergent.

**Extended Data Table 1.** CryoEM data, structure refinement, and validation. Note that CXCR4EM/REGN7663 Fab trimer and tetramer structures were obtained from the same dataset.

|  | CXCR4<sub>EM</sub>/<br>G<sub>i</sub> | CXCR4<sub>EM</sub>/<br>G<sub>i</sub>/CXCL12 | CXCR4<sub>EM</sub>/<br>G<sub>i</sub>/<br>AMD3100 | CXCR4<sub>EM</sub>/<br>G<sub>i</sub>/<br>REGN7663 Fab | CXCR4<sub>EM</sub>/<br>REGN7663<br>Fab | CXCR4<sub>EM</sub>/<br>REGN7663<br>Fab Trimer | CXCR4<sub>EM</sub>/<br>REGN7663<br>Fab Tetramer |
| --- | --- | --- | --- | --- | --- | --- | --- |
| **Data collection and processing** |  |  |  |  |  |  |  |
| Magnification | 165,000 | 105,000 | 105,000 | 105,000 | 105,000 | 105,000 | 105,000 |
| Voltage (kV) | 200 | 300 | 300 | 300 | 300 | 300 | 300 |
| Electron exposure (e<sup>-</sup>/Å<sup>2</sup>) | ~50 | ~40 | ~40 | ~40 | ~40 | ~40 | ~40 |
| Defocus range | -1.0 to -2.0 | -1.2 to -2.2 | -1.2 to -2.2 | -1.2 to -2.2 | -1.2 to -2.2 | -1.2 to -2.2 | -1.2 to -2.2 |
| Pixel size (Å) | 0.696 | 0.85 | 0.85 | 0.85 | 0.85 | 0.85 | 0.85 |
| # of Movies | 9,693 | 11,922 | 10,013 | 10,346 | 5,827 | 5,329 | 5,329 |
| Initial number of particles | 5.0M | 13.0M | 8.1M | 7.1M | 6.2M | 3.0M | 3.0M |
| Particles selected after 2D classification | 615K | 805K | 803K | 1.03M | 639K | 585K | 585K |
| Final selected particles | 183,399 | 87,963 | 193,216 | 148,703 | 102,810 | 27,104 | 31,775 |
| Symmetry imposed | C1 | C1 | C1 | C1 | C1 | C3 | C4 |
| Map resolution (Å) | 2.72 | 3.29 | 3.15 | 3.36 | 3.10 | 3.35 | 3.38 |
| FSC threshold | 0.143 | 0.143 | 0.143 | 0.143 | 0.143 | 0.143 | 0.143 |
| Refinement |  |  |  |  |  |  |  |
| Initial Model used | This study | 3HP3, this study | This study | 4RWS, 7T2G | This study | This study | This study |
| **Model composition** |  |  |  |  |  |  |  |
| Non-hydrogen atoms | 6,963 | 7,401 | 7,006 | 10,409 | 4,107 | 12,684 | 16,968 |
| Protein residues | 889 | 966 | 892 | 1,339 | 513 | 1,557 | 2,056 |
| Ligands | 1 | 1 | 2 | 1 | 1 | 9 | 20 |
| **R.m.s. deviations** |  |  |  |  |  |  |  |
| Bond lengths (Å) | 0.005 | 0.002 | 0.002 | 0.005 | 0.002 | 0.003 | 0.004 |
| Bond angles (°) | 0.653 | 0.493 | 0.457 | 0.642 | 0.629 | 0.715 | 0.700 |
| **Validation** |  |  |  |  |  |  |  |
| MolProbity score | 1.74 | 1.56 | 1.41 | 1.69 | 1.50 | 1.55 | 1.75 |
| Rotamer outliers (%) | 0.00 | 0.00 | 0.00 | 0.00 | 0.00 | 0.67 | 0.00 |
| Clash score | 8.65 | 5.56 | 4.24 | 6.54 | 4.99 | 7.06 | 8.53 |
| Ramachandran plot |  |  |  |  |  |  |  |
| Favored (%) | 96.02 | 96.12 | 96.71 | 95.31 | 96.42 | 97.08 | 95.85 |
| Allowed (%) | 3.98 | 3.88 | 3.29 | 4.46 | 3.58 | 2.92 | 4.15 |
| Disallowed (%) | 0.00 | 0.00 | 0.00 | 0.23 | 0.00 | 0.00 | 0.00 |
| **Deposition ID** |  |  |  |  |  |  |  |
| PDB | 8U4N | 8U4O | 8U4P | 8U4Q | 8U4R | 8U4S | 8U4T |
| EMDB | 41888 | 41889 | 41890 | 41891 | 41892 | 41893 | 41894 |

